# Evolutionary consequences of repeated loss of distyly in *Linum*

**DOI:** 10.64898/2026.03.03.709227

**Authors:** Zoé Postel, Panagiotis-Ioannis Zervakis, Marco Fracassetti, Aleksandra Losvik, Matias Wanntorp, Lucile Soler, Allison Churcher, Aelys M. Humphreys, Tanja Slotte

**Affiliations:** Department of Ecology, Environment and Plant Sciences, Science for Life Laboratory, Stockholm University, Stockholm, Sweden; SciLifeLab, National Bioinformatics Infrastructure Sweden (NBIS), Department of Medical Biochemistry and Microbiology, Uppsala University, Uppsala, Sweden; Department of Plant Physiology, National Bioinformatics Infrastructure Sweden, Science for Life Laboratory, Umeå University, Umeå, Sweden

**Keywords:** <u>Key words</u>: mating system shift, mixed-mating, selfing, chloroplast genome, selective pressure

## Abstract

The breakdown of distyly, a polymorphism that promotes disassortative pollination between two floral morphs, is often assumed to increase self-fertilization, reducing the efficacy of selection and challenging long-term species persistence. We tested whether repeated losses of distyly *Linum* were associated with genomic signatures of selfing by 1) testing for relaxed selective pressure in homostylous relative to distylous lineages, and 2) by characterising population genomic patterns of the homostylous *Linum leonii* in comparison to its distylous close relative *Linum perenne*. We generated whole-genome sequences and target-capture data from sixteen *Linum* species, and additionally built a high-quality genome assembly and acquired population-level whole-genome sequencing data for *L. leonii* (n=20). We reconstructed plastome phylogenies, estimated selective pressure for chloroplast and nuclear genes, inferred ancestral floral morph states, and tested for signatures of selfing in homostylous lineages. Compared to theoretical expectations, results were mixed, with partial identification of relaxed selective pressure in homostyles. Population genomic analyses of *L. leonii* revealed a moderate selfing rate of 0.32, suggesting that loss of distyly was associated with mixed mating rather than selfing, contrary to previous results on loss of distyly. Reduced nucleotide diversity and evidence for relaxed selection efficacy in *L. leonii* were instead consistent with a historical bottleneck. In *Linum,* the genomic consequences are more heterogeneous than generally assumed, and likely depend on species-specific evo-demographic history. This study highlights the complex evolutionary dynamics associated with the breakdown of distyly and emphasizes the need for comparative population genomic studies to clarify how such transitions shape evolutionary processes.

**Significance statement:** Plant mating system variation is central to evolution as it shapes genetic diversity, adaptability and fitness. Loss of distyly, an iconic example of a complex mating system favouring cross-pollination, can drive shifts from outcrossing to selfing, with potentially severe evolutionary consequences for the long-term persistence of the species in which it occurs. Using high-quality genome assembly and omic data for multiple *Linum* species, we tested for relaxed selective pressure in homostylous compared to distylous lineages, and tested for a population genomic signature of selfing in homostylous *Linum leonii* compared to the closely related distylous *Linum perenne*. Contrary to theoretical expectations, evidence for relaxation of selection was mixed in *Linum* homostyles and *L. leonii* did not exhibit a genomic signature of selfing. Our study reveals multiple evolutionary pathways following the loss of distyly, and highlights how mating system transitions, together with complex demographic processes, shape plant genetic diversity and evolution.

## Introduction

Flowering plants display a remarkable array of intricate floral adaptations to attract pollinators and promote efficient outcrossing. Among these, the floral polymorphism distyly is a prominent example. In distylous species, populations comprise two floral morphs: the L-morph (“pin”, long-styled, short anther filaments) and the S-morph (“thrum”, short-styled, long anther filaments) (Figure 1-A). The reciprocal positioning of sexual organs enhances intermorph pollen transfer between morphs by pollinators (Simón-Porcar et al. 2024), usually coupled with a heteromorphic self-incompatibility (SI) mechanism which limits self- and intra-morph fertilization (Darwin 1877; Charlesworth and Charlesworth 1979; Lloyd and Webb 1992; Barrett 2019). This complex polymorphism thus promotes efficient outcrossing and inbreeding avoidance.

**Figure 1.**
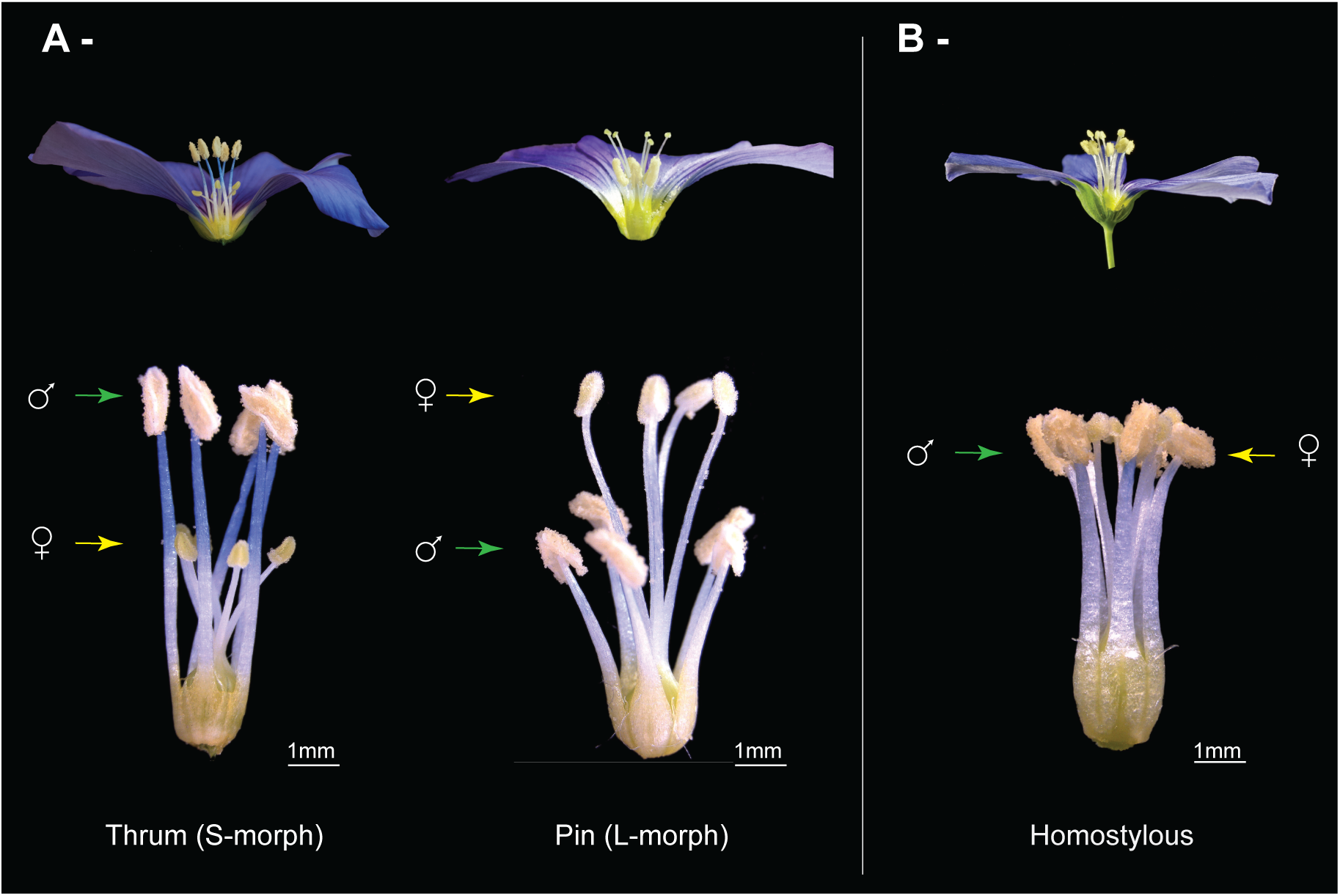
Photographs of the flowers of A) distylous *L. perenne,* with the thrum (S-morph) morph to the left and pin morph (L-morph) to the right and B) *L. leonii* (homostylous). In the top panel, petals and sepals have been removed from one side of the flower to better visualize the positions of the sexual organs. In the bottom panel, all petals and sepals have been removed. For both species, arrows indicate the positions of anthers (green) and stigmas (yellow). Scale bars (1 mm) are shown for the bottom panel.

Although distyly provides adaptive advantages by promoting outcrossing and reducing inbreeding depression, it has repeatedly been lost across angiosperms (Barrett 2019). Theoretically, multiple evolutionary trajectories may follow the breakdown of distyly, each with distinct effects on genetic diversity, morph frequencies, and mating patterns (Ganders 1979; Yuan et al. 2017; Barrett 2019; Yuan et al. 2023). If loss of SI occurs while pollinators still confer intermorph pollen transfer, populations may evolve self-compatible (SC) distyly, retaining disassortative mating but allowing intramorph crosses and limited selfing. Alternatively, when pollinators promote assortative mating, monomorphic SC outcrossing or mixed-mating populations with reduced herkogamy may prevail. The derived state of floral monomorphism with anthers and stigmas at the same height and absence of herkogamy is termed homostyly (Darwin 1877; Ganders 1979). Finally, in environments poor in pollinators, selection for reproductive assurance favours the evolution of SC and homostylous flowers (Figure 1-B) which enables efficient autonomous self-fertilization (selfing) (Lloyd and Webb 1992; Yuan et al. 2023). While homostylous species are usually SC, exceptions exist, e.g. in Oleaceae where derived floral homomorphism and diallelic SI co-occurs in some species (Castric et al. 2024; Raimondeau et al. 2024).

Of these three main evolutionary pathways, in most of the systems studied so far, loss of distyly was associated with the evolution of homostyly (Ganders 1979; Barrett 2019, Figure 1-B) and a shift to higher selfing rate (Yuan et al. 2017; Zhong et al. 2019; Mora-Carrera et al. 2021; Gutiérrez-Valencia, P.-I. Zervakis, et al. 2024). Little is known about how frequently loss of distyly results in other possible outcomes involving SC and intermediate outcrossing levels (but see (Barrett and Shore 2008; de Vos et al. 2018; Yuan et al. 2019)).

Considering homostyly as the evolutionary pathway most commonly documented in cases of distyly breakdown, several evolutionary consequences are expected (Charlesworth and Charlesworth 1979). Typically, selfing species exhibit a ‘selfing syndrome’: they have smaller flowers with lower pollen-to-ovule ratios and reduced nectar and scent, reflecting the combined effects of altered selective pressures related to pollinator attraction and autonomous selfing ability (Darwin 1876; Sicard and Lenhard 2011; Cutter 2019; Tsuchimatsu and Fujii 2022). Shifts to high selfing rates also leave genomic signatures, defined as the ‘genomic selfing syndrome’ (reviewed in Wright et al. 2013; Slotte 2014; Cutter 2019; Wang et al. 2021). Such a syndrome entails a decrease in the effective population size (*N*_*e*_) and in the effective recombination rate. Together, this results in a decline in genetic diversity and impacts the efficacy of selection (Burgarella and Glémin 2017; Hartfield et al. 2017). If the shift to selfing is relatively recent, selection efficacy could increase as increased homozygosity due to selfing would help reveal, previously masked heterozygous recessive deleterious mutations, resulting in purging (Arunkumar et al. 2015; Burgarella and Glémin 2017). But if the shift occurred further back in time, an increase in selfing rate could lead to a decrease in selection efficacy and accumulation of mildly and slightly deleterious mutations (Burgarella and Glémin 2017). Therefore, all else being equal, a decrease in the efficacy of purifying selection should yield a higher ratio of non-synonymous (N) to synonymous (S) mutations (π_*N*_⁄π_*S*_ if based on polymorphism data or *d*_*N*_⁄*d*_*S*_ if estimated with divergence data) for homostylous species than their outcrossing distylous relatives.

Most previous studies on the impact of selfing on selection have looked for a decrease in selection efficacy. Polymorphism-based estimates of selection efficacy are consistent with theoretical expectations (Slotte et al. 2010; Arunkumar et al. 2015; Douglas et al. 2015; Chen et al. 2017; Muyle et al. 2021) but divergence data show mixed support (Escobar et al. 2010; Glémin and Muyle 2014; Wang et al. 2021). While genomic signatures of selfing have primarily been documented in population genomic contrasts of pairs of selfing and outcrossing species, genus-wide comparative genomic assessments across multiple shifts to selfing (e.g. Wang et al. 2021) are needed to draw general conclusions on the genomic and evolutionary impact of plant mating system shifts (Mattila et al. 2020). With a few exceptions (Glémin and Muyle 2014; Wang et al. 2021), most earlier studies focused on the selective pressure at the nuclear level and rarely studied selection on organellar genomes. Yet, analyzing organellar genes alongside nuclear loci helps clarify differences in selective pressure between distylous and homostylous species. Because organellar genomes are haploid and uniparentally inherited, they are unaffected by dominance, unlike nuclear genes where recessive deleterious mutations may be masked in outcrossing and exposed under selfing, potentially increasing purifying selection. Nuclear genes are also more strongly influenced by linkage and hitchhiking, whereas organellar genes are only indirectly affected by genome-wide linkage due to selfing (Glémin & Muyle 2014). Comparing these genomes therefore reveals whether differences in selective pressure between mating systems are consistent across contrasting dominance and population genetic contexts.

*Linum* (flax) is a valuable system for studying both the evolution and breakdown of distyly. The genus displays remarkable stylar polymorphism including distyly, stigma-height dimorphism, and recurrent transition to homostyly across distantly related species (i.e., multiple times independently) (Ruiz-Martín et al. 2018; Maguilla et al. 2021). In *Linum*, distylous and style polymorphic species typically have heteromorphic SI (e.g., *Linum perenne, Linum grandiflorum*: (Darwin 1863; Darwin 1877; Murray 1986; McDill et al. 2009), *Linum flavum*, *Linum tenue*, *Linum austriacum, Linum narbonense, Linum hirsutum*: (Murray 1986)*, Linum suffruticosum* s.l.: (Nicholls 1985)), whereas homostylous species are typically SC (e.g., *Linum bienne*: (Darwin 1863; Darwin 1877); *Linum trigynum*: (Murray 1986; Gutiérrez-Valencia, et al. 2024) ; *Linum lewisii*: (Cockerell 1902), *Linum tenuifolium* s.l.: (Nicholls 1985; McDill et al. 2009), *Linum leonii*: (Ockendon 1968)). Earlier work revealed reduced nucleotide diversity, elevated inbreeding coefficients, reduced efficacy of selection, and increased population structure in the annual homostylous SC *L. trigynum*, compared to the closely related distylous SI *L. tenue*. Results for *L. trigynum* provide an example of the genomic selfing syndrome (Cutter 2019) following transition to high selfing rates (Gutiérrez-Valencia, P.-I. Zervakis, et al. 2024).

Here, we leverage the repeated transitions from distyly to homostyly in *Linum* and generate new genetic resources to assess whether the loss of distyly is consistently associated with a shift to a higher selfing rate in this genus. Specifically, we address two overarching questions: (i) Do homostylous species across *Linum* systematically exhibit evidence of whole-genome relaxed selection, as expected when loss of distyly leads to an increase in selfing rate? (ii) By focusing on *Linum leonii* (section *Linum*), a homostylous SC blue flax (Ockendon 1968), do we find similar support for a genomic selfing syndrome as in *L. trigynum* (section *Linopsis*), a distantly related homostylous species (McDill et al. 2009; Gutiérrez-Valencia et al. 2024)? To assess selection across a broad phylogenetic sample of *Linum*, we used whole-genome and target-capture sequencing of nineteen species (13 distylous and 6 homostylous). For 14 out of the 19 *Linum* species analyzed here, SI/SC status is known, and in all cases distylous species are SI and homostylous species SC (e.g., (Nicholls 1985; Murray 1986; Dulberger 1987; Kearns and Inouye 1994; Gutiérrez-Valencia et al. 2024)). Moreover, we sequenced, assembled and annotated a high-quality genome of *L. leonii* and used population genomic analyses to test for a genomic selfing syndrome through quantification of genome-wide nucleotide diversity, inbreeding coefficients, and assessment of the efficacy of purifying and positive selection. We compared its genomic signatures to those of the closely related SI distylous blue flax, *L. perenne*, for which a high quality genome assembly as well as population genomic data were already available . By integrating genomic and population-genetic analyses across *Linum* in a comparative framework, our study uncovers how alternative evolutionary pathways from distyly to homostyly and selfing may shape genomic diversity and selection efficacy. The results contribute to an improved understanding of evolutionary transitions in mating systems and how they impact genome evolution over both shorter and longer time scales.

## RESULTS

### Testing for relaxed selection in homostylous Linum species

#### Phylogeny reconstruction

We generated whole-genome sequencing short-read data for sixteen *Linum* species and downloaded data from NCBI for three additional species and an outgroup, *Tirpitzia sinensis* (Suppl. Table S1). Chloroplast genomes were reconstructed using GetOrganelle (Jin et al. 2020). Overall, the chloroplast genome assemblies were of good quality, with deep coverage of 490.45x and an average length of 157 405 base pairs (bp) (Suppl. Table S2). With GeSeq annotations (Tillich et al. 2017), we retrieved the four genomic parts usually composing chloroplast genomes: large single-copy (LSC), small single-copy (SSC), and two inverted-repeat regions (IR) (Suppl. Figure S1). Out of the 115 genes present in the database, 88% (i.e., 101/115) were suitable for further analysis (see Methods for details). We then restricted the dataset to protein-coding genes (i.e., 67/101) and constructed a concatenation of these aligned gene sequences, with a total length of 48 513 bp.

This concatenation was then used to reconstruct a phylogenetic tree using RAxML (Stamatakis 2014) with a GTR-GAMMA model of nucleotide substitution. The resulting phylogeny was well supported, with all nodes supported by bootstrap values of 100%, except for one node comprising *L. campanulatum, L. flavum, L. capitatum* and *L. thracicum*, that had a bootstrap value around 30% (Figure 2). All nineteen *Linum* species fell into two major clades, which correspond to clade A (section *Linum/Dasylinum*) and clade B2 (section *Linopsis/Syllinum*) found previously (McDill et al. 2009; Ruiz-Martín et al. 2018; Maguilla et al. 2021). Clade A comprised four homostylous and five distylous species and clade B2 eight distylous and two homostylous species. Clade A branches were on average longer than those of clade B2 (Figure 2).

**Figure 2.**
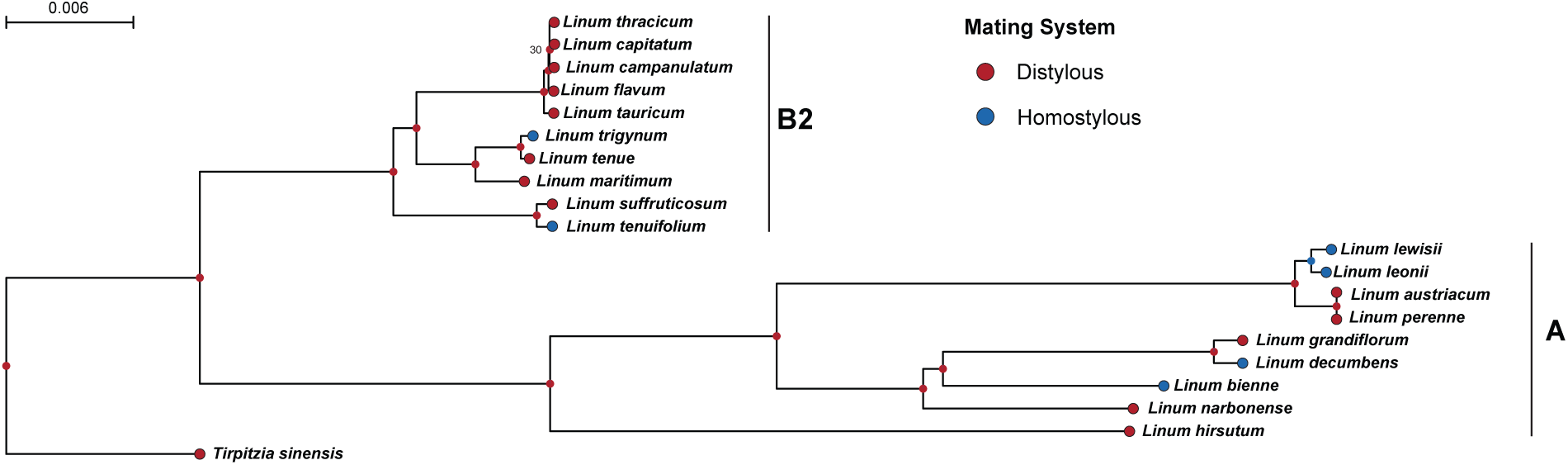
Chloroplast phylogeny and ancestral state reconstruction. The chloroplast phylogenetic tree was reconstructed using RAxML and a GTR-GAMMA model of nucleotide substitution. Node support values are shown where bootstrap support was <100% (only for one node). Ancestral state reconstruction of mating system (distylous/ homostylous) was performed on the time-calibrated chloroplast phylogeny using the Hidden rates model (corHMM). Ancestral states were simulated in plotSimmap. Node and tip colors indicate distylous (red) and homostylous (blue) states.

To assess the selective pressure on the nuclear genomes, we generated target capture data for the same sixteen *Linum* species, using the Angiosperms353 probe set (Johnson et al. 2019). Gene alignments were retrieved using Hybpiper (Johnson et al. 2016) and T-coffee, further selecting only the 129 genes with good quality alignments (T-Coffee alignment score > 80 and no internal stop codon) and gene sequences for all twenty species. After building 129 gene trees with RAxML, we ran ASTRAL (Zhang et al. 2018) to get a nuclear-based phylogeny and check for congruence between trees. The nuclear-based ASTRAL phylogeny had the same topology as the chloroplast phylogeny, exhibiting similarly high node support (i.e. bootstrap value >80). The only discrepancy between phylogenies was for the same group of species with a lower node support in the chloroplast tree, which formed one clade in the ASTRAL phylogeny, with all internal nodes exhibiting bootstrap values below 80 (Suppl. Figure S2). This discrepancy between chloroplast and nuclear trees indicates that there probably is not enough variation to resolve this clade (either its existence – based on high similarity among chloroplast DNA sequences of these four species – or its resolution, based on the nuclear sequences). This could also reflect incomplete lineage sorting, given the recent divergence suggested by the short internal branches in this clade. Past hybridization events can also not be excluded, although the lack of strong, well-supported topological conflict between nuclear and chloroplast trees suggest that limited phylogenetic signal is likely the primary explanation.

#### Ancestral state reconstruction

To infer whether internal nodes were distylous or homostylous, we performed ancestral state reconstruction using the chloroplast tree. After time-calibrating the tree, we assessed ancestral states by designing custom transition rate matrices using corHMM (Boyko and Beaulieu 2021) in R. The best-fitting model was the one with a transition rate from distyly to homostyly = 0.01 transitions/My and from homostyly to distyly = 1 × 10^-4^ transitions/My (negative log-likelihood: -28.89; sample-size adjusted Akaike information criterion (AICc) = 61.77). The best-fitting model suggested five independent transitions from distyly to homostyly across the six homostylous species analysed and no reversals to distyly (Figure 2). Overall, most deep internal nodes were reconstructed as distylous with high support (marginal probability (*Pr*) > 0.99) with only one node (the common ancestor of *L. leonii* and *L. lewisii*) inferred as being ancestrally homostylous (*Pr* ≈ 0.97).

#### Divergence-based analyses of selection

To test for relaxed selective pressure in homostylous species of *Linum*, we estimated *d*_*N*_⁄*d*_*S*_ values using PAML – codeml (Yang 2007) and different branch-models on chloroplast and nuclear genes (Suppl. Table S3). This method allows *d*_*N*_⁄*d*_*S*_ to vary along annotated phylogenetic branches and the best branch-model is selected using a log likelihood-ratio test. For the nuclear gene sets, among the 129 genes used for the ASTRAL phylogeny, 50 had a paralog warning from Hybpiper. We thus ran the PAML analysis on two nuclear gene sets: the 79 genes without this warning (later referred to as the D1 dataset) and the full set of 129 genes (later referred as the D2 dataset). PAML was run separately on concatenations of D1, D2 and chloroplast genes, respectively. In the main text we are detailing results for D2 while results for D1 can be found in the supplementary.

For both concatenated chloroplast gene and nuclear D2 gene sets, the best branch-model was H2 (i.e., one *d*_*N*_⁄*d*_*S*_ estimated per mating system in each clade) (Table 1). Overall, regarding chloroplast genes, clade A lineages exhibited slightly higher *d*_*N*_⁄*d*_*S*_ compared to clade B2 (mean *d*_*N*_⁄*d*_*S*_ clade A = 0.19 vs clade B2 = 0.11) (Table 2). Homostylous lineages of clade B2 exhibited higher *d*_*N*_⁄*d*_*S*_ values compared to distylous lineages (Table 2). For clade A, *d*_*N*_⁄*d*_*S*_ values were equivalent between homostylous and distylous lineages (Table 2). For D2 nuclear genes, we observed higher *d*_*N*_⁄*d*_*S*_ values for homostylous lineages of clade A and no differences between distylous/homostylous lineages of clade B2 (Table 2). Results were similar for D1 nuclear genes, with clade A homostylous lineages exhibiting higher *d*_*N*_⁄*d*_*S*_ values than the distylous lineages and similar estimates for both distylous and homostylous lineages of clade B2 (Suppl. Table S4 and S5).

**Table 1.**
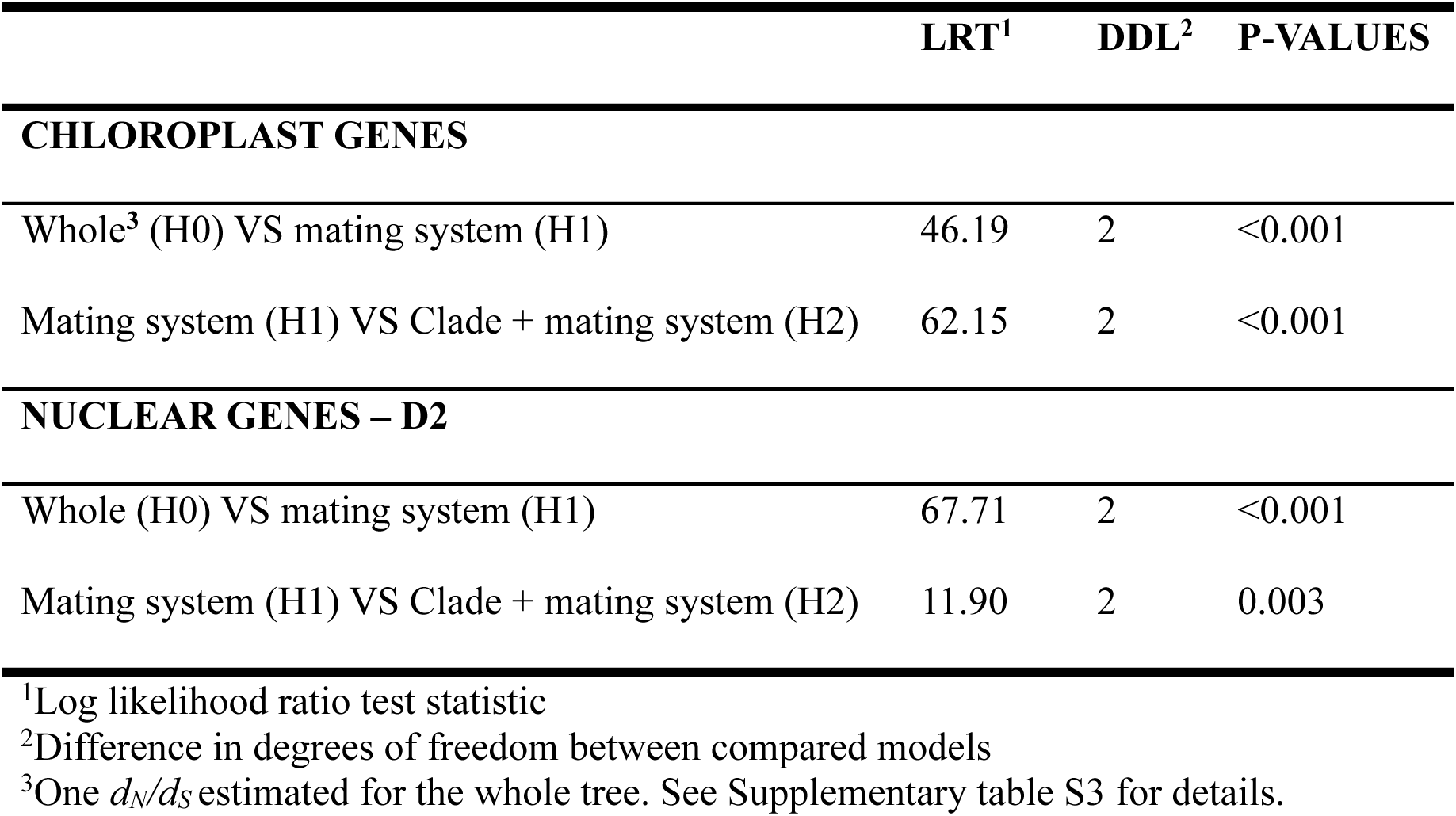
Log likelihood-ratio test results for chloroplast and nuclear genes (D2 data set). The table shows results of comparisons of models with one *d_N_/d_S_* value (H0) vs separate values for distylous and homostylous lineages (H1), and with clade as an added factor (H2).

|  | LRT<sup>1</sup> | DDL<sup>2</sup> | P-VALUES |
| --- | --- | --- | --- |
| **CHLOROPLAST GENES** |  |  |  |
| Whole<sup>3</sup> (H0) VS mating system (H1) | 46.19 | 2 | <0.001 |
| Mating system (H1) VS Clade + mating system (H2) | 62.15 | 2 | <0.001 |
| **NUCLEAR GENES – D2** |  |  |  |
| Whole (H0) VS mating system (H1) | 67.71 | 2 | <0.001 |
| Mating system (H1) VS Clade + mating system (H2) | 11.90 | 2 | 0.003 |

<sup>1</sup>Log likelihood ratio test statistic
<sup>2</sup>Difference in degrees of freedom between compared models
<sup>3</sup>One $d_N/d_S$ estimated for the whole tree. See Supplementary table S3 for details.

**Table 2.** Estimates of *d*_*N*_⁄*d*_*S*_ from PAML under the codeml H1 and H2 branch-model for chloroplast and nuclear genes (D2 dataset). Standard errors are indicated after each estimate.

| Clade | Hypothesis | Branch model Categories | Chloroplast genes | Nuclear genes |
| --- | --- | --- | --- | --- |
| Clade A | H1 | All clade | $0.19 \pm 0.006$ | $0.10 \pm 0.004$ |
| | H2 | Distylous | $0.19 \pm 0.006$ | $0.09 \pm 0.002$ |
| | | Homostylous | $0.18 \pm 0.02$ | $0.12 \pm 0.005$ |
| Clade B2 | H1 | All clade | $0.11 \pm 0.006$ | $0.10 \pm 0.002$ |
| | H2 | Distylous | $0.11 \pm 0.006$ | $0.10 \pm 0.002$ |
| | | Homostylous | $0.17 \pm 0.05$ | $0.09 \pm 0.008$ |

To check whether the elevated *d*_*N*_⁄*d*_*S*_ for some lineages was the results of increased *d*_*N*_ and not reduced *d*_*S*_, we extracted *d*_*N*_ and *d*_*S*_ estimates for each external branch of the best-fitting branch-model in PAML. We then computed mean values per clade and per mating system within clade and tested for statistical significance. Overall, neither *d*_*S*_ nor *d*_*N*_ differed significantly between clades or between mating system within clades, suggesting that only slight differences were detected between clades and mating system in both genomic compartments (for the nuclear dataset: *d*_*S*_ Kruskal-Wallis test chi2 (KW chi2) = 3.44, p-value = 0.33 & *d*_*N*_ KW chi2 = 4.47, p-value = 0.22 ; for the chloroplast dataset: *d*_*S*_ Kruskal-Wallis test chi2 (KW chi2) = 2.25, p-value = 0.52 & *d*_*N*_ KW chi2 = 3.17, p-value = 0.37). Estimates of *d*_*S*_ were slightly higher in clade A than in clade B2 for both chloroplast and nuclear genes (i.e. clade A chloroplast = 1.5 × 10^−3^ vs clade B2 chloroplast = 8.9 × 10^−4^ and nuclear = 0.04 vs 0.02, respectively; Table 3). In contrast, *d*_*N*_ estimates were similar between clade A and clade B2, for both genomic compartments. With respect to mating system, homostylous species exhibited marginally higher *d*_*S*_for chloroplast genes in both clades but this pattern was observed only in clade B2 for the nuclear genes (0.03 for the homostylous vs 0.02 for the distylous) (Table 3). For *d*_*N*_, homostylous species in clade B2 – but not clade A – exhibited slightly higher estimates for the chloroplast genes (7.8 × 10^−5^ for the distylous vs 3.0 × 10^−4^ for the homostylous) whereas no differences were detected the nuclear genes (Table 3).

**Table 3.** Estimates of mean *d*_*N*_ and *d*_*S*_ per clade and mating system from PAML under the codeml H2 branch-model, for chloroplast and nuclear D2 datasets.

| Statistic | Clade | Mating system | Divergence |  |
| --- | --- | --- | --- | --- |
|  |  |  | Chloroplast | Nuclear, D2 |
| $d_S$ | Clade A | All | $1.5 \times 10^{-3}$ | 0.04 |
| | | Distylous | $6.2 \times 10^{-4}$ | 0.04 |
| | | Homostylous | $4.3 \times 10^{-3}$ | 0.04 |
| | Clade B2 | All | $8.9 \times 10^{-4}$ | 0.02 |
| | | Distylous | $7.1 \times 10^{-4}$ | 0.02 |
| | | Homostylous | $1.8 \times 10^{-3}$ | 0.03 |
| $d_N$ | Clade A | All | $2.9 \times 10^{-4}$ | $4.4 \times 10^{-3}$ |
| | | Distylous | $1.3 \times 10^{-4}$ | $3.9 \times 10^{-3}$ |
| | | Homostylous | $7.7 \times 10^{-4}$ | $5.0 \times 10^{-3}$ |
| | Clade B2 | All | $1.1 \times 10^{-4}$ | $2.0 \times 10^{-3}$ |
| | | Distylous | $7.8 \times 10^{-5}$ | $1.8 \times 10^{-3}$ |
| | | Homostylous | $3.0 \times 10^{-4}$ | $3.0 \times 10^{-3}$ |

### Characterizing the evolutionary consequence of loss of distyly in L. leonii

#### High-quality genome assembly of L. leonii

To test for a selfing syndrome in the homostylous *L. leonii*, we first built a high-quality genome assembly. We generated a genome assembly for one individual of *L. leonii* using PacBio HiFi and Hi-C data (Suppl. Table S6). The resulting assembly exhibited a BUSCO score of 93.9% indicating good completeness. The N50 was 21.08 Mb, highlighting assembly contiguity, and the total length of the assembly was similar to genome size estimated based on flow cytometry (Suppl. Table S6). The primary assembly was annotated with ab-initio and transcriptomic data and led to the identification of c. 45 000 protein-coding genes (Dataset S1). It is likely that the high gene number in *L. leonii* is the results of an ancient whole-genome duplication in the clade (Sveinsson et al. 2014). In total, 74,75% of the genome was annotated as repetitive elements, with the majority being LTR retroelements (55.3%) and DNA transposons (11.17%) (Suppl. Table S7).

#### Population structure and time of split between L. perenne and L. leonii

To conduct population genomic analyses, we acquired population genomic data for twenty individuals of *L. leonii* and additionally used assembly and population genomic data of three natural populations of the closely related distylous *L. perenne* (Zervakis, Postel et al. 2025) (ENA: PRJEB88074, Suppl. Table S8). Variants were called and filtered after mapping all reads to (1) the *L. perenne* genome assembly and (2) to each assembly separately (see Methods and Suppl. Table S9 for details).

As the two species have partially overlapping geographical distributions (Suppl. Fig. S3) and hybridization could confound analyses of loss of distyly in *L. leonii*, we first determined population differentiation and the degree of gene flow between these species. For this purpose we conducted population structure analyses using Principal Component Analysis (PCA) in PLINK (Chang et al. 2015) and ran ADMIXTURE (Alexander et al. 2009) on the VCF (Variant Call Filtering) file with all reads mapped to *L. perenne.* In the PCA analysis, the first principal component explained 36.9% of the variation and clearly separated *L. leonii* from *L. perenne*, whereas the second and third principal components (explaining 10.3% vs 6.5% of the variation, respectively) reflected population structure within *L. perenne* (Suppl. Figure S4). ADMIXTURE results for K = 8, which had the lowest cross-validation score, were in broad agreement with PCA results, showing clear separation of *L. leonii* from *L. perenne* (Figure 3-A, Suppl. Table S10). This suggests that there is very limited gene flow between *L. leonii* and *L. perenne*.

**Figure 3.**
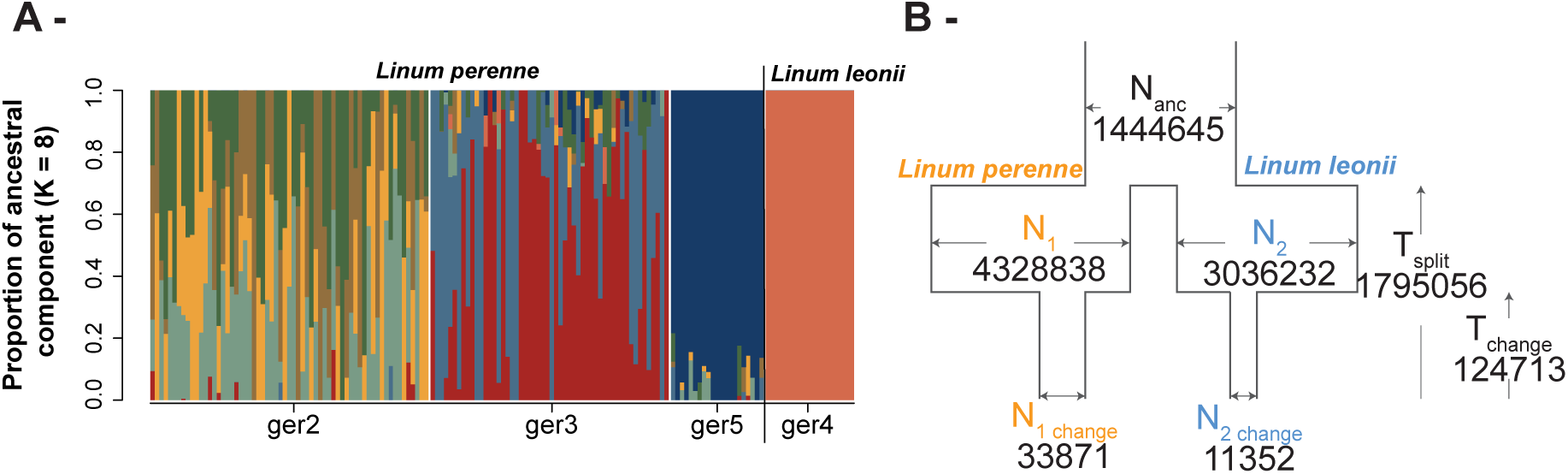
Population structure and demographic history of *L. perenne* and *L. leonii*. **A)** ADMIXTURE results with K = 8. The y-axis indicates the proportion of ancestry assigned to each ancestral component K, whereas the x-axis corresponds to individuals from different populations of *L. perenne* and *L. leonii*. The choice of K value was based on cross-validation error values. Results highlight clear population structure between the two species. **B)** Inferred demographic history model between *L. leonii* and *L. perenne* population “ger2”. Both species experienced an increase and subsequently a decrease in effective population size after their split. Estimated effective population sizes (individuals) are given, as well as estimates of split times and the length of the bottleneck (in years). N_anc_: effective population size of the ancestral population. N_1,2_: the effective population size of *L. perenne* and *L. leonii* respectively, immediately after the split. N_1,2 change_: the bottlenecked effective population size of *L. perenne* and *L. leonii* respectively. T_split_: time since the species split. T_change_:onset of the bottleneck.

To estimate the timing of the split, which is also the upper limit for the shift to SC in *L. leonii*, we modelled demographic history in pairwise comparisons between *L. perenne* and *L. leonii* using the same VCF and *dadi* models with inbreeding (Gutenkunst et al. 2009; Blischak et al. 2020). The species split was inferred at the earliest ∼2 Mya (*L. perenne* ger2, ger5 and *L. leonii*) and latest at ∼1 Mya (*L. perenne* ger3 and *L. leonii*), and all best-fit pairwise split models included population size change following the split (Suppl. Table S11). The best-fitting model between *L. perenne* population ger2 and *L. leonii* (i.e. *“no_mig_size”*, Suppl. Fig. S5) indicated that both populations experienced an initial population growth after their split, followed by a decline in effective population sizes at ∼125 kya: from 1,400,000 in the common ancestor to around 60,000 in *L. perenne* ger2 and 10,000 in *L. leonii* (Figure 3-B, Suppl. Table S11). The inbreeding coefficient estimated by *dadi* was close to the values obtained with *F_IS_* (*F_ger2_* = 0.01, *F_leonii-ger2_* = 0.20, Figure 4-A, see per-population *F_IS_*comparisons below). For the two other comparison (i.e. *L. leonii* vs *L. perenne* – ger3 and *L. leonii* vs *L. perenne* – ger5), similar tendencies of *N*_*e*_ decline compared to the common ancestor and low to medium inbreeding coefficients were inferred (Suppl. Table S11). Further details for these can be found in Suppl. Note S1 and Suppl. Table S11.

**Figure 4.**
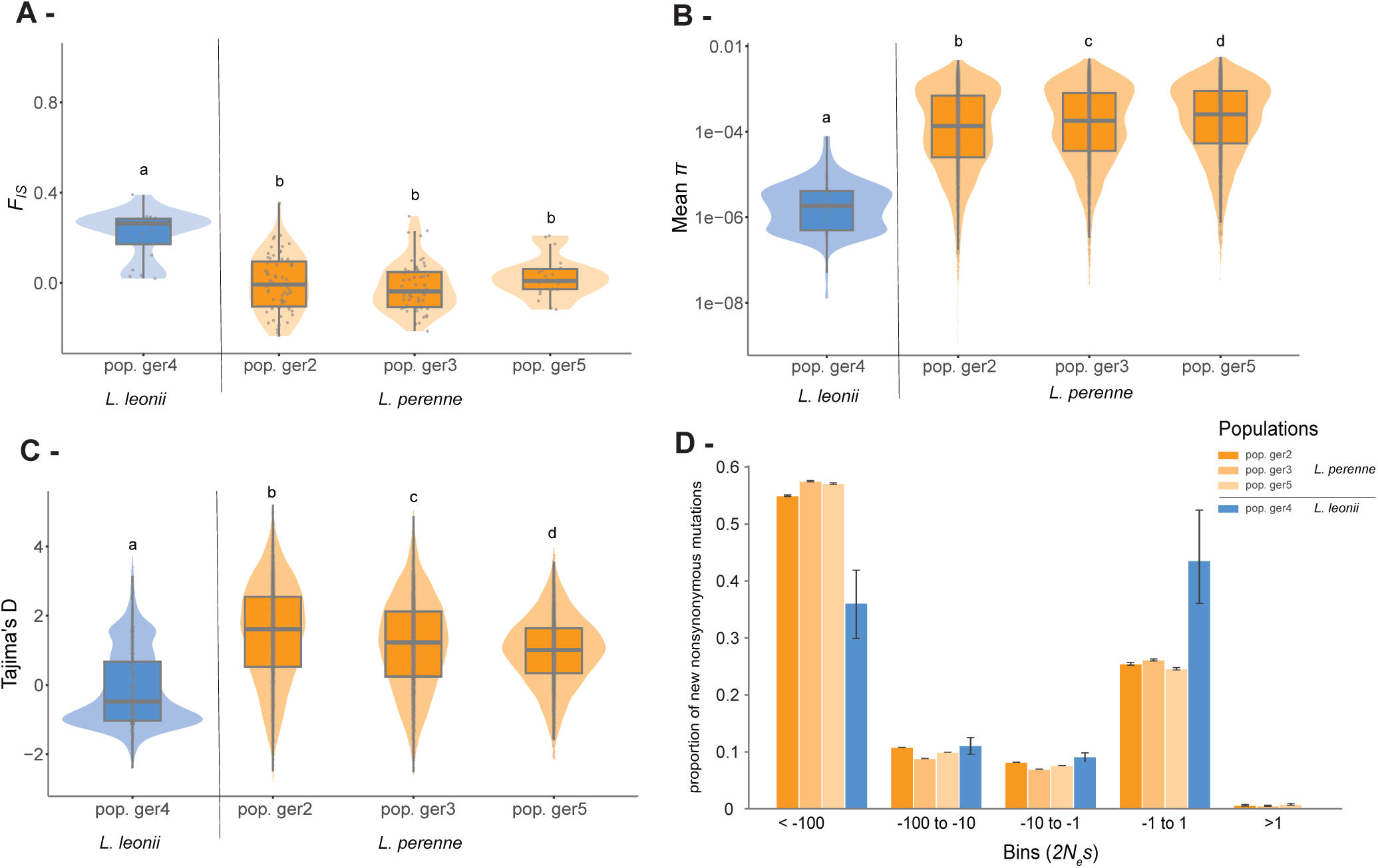
Population genomic statistics for *L. leonii* and *L. perenne*. **A)** Whole-genome estimates of mean *F_IS_*. Different letters indicate significant differences between the population estimates (Kruskal-Wallis chi-squared = 29.52, df = 3, p-value < 0.001 followed by Dunn-test with Benjamini-Hochberg p-value adjustment). *L. leonii* exhibited significantly higher values of *F_IS_* compared to *L. perenne*, yet well below 1, suggesting that *L. leonii* might be mixed-mating rather than predominantly selfing. **B)** Whole-genome estimates of mean π by 100kb window. Differences were statistically significant for all pairs of populations (Kruskal-Wallis chi-squared = 514, df = 3, p-value < 0.001 followed by Dunn-test with Benjamini-Hochberg p-value adjustment). *L. leonii* exhibited significantly lower values of π highlighting whole-genome decrease in genetic diversity as expected when shifting to higher selfing rate or after a recent bottleneck. **C)** Whole-genome estimates of mean Tajima’s D by 100kb window. Differences were statistically significant for all pairs of populations (Kruskal-Wallis chi-squared = 2176.50, df = 3, p-value < 0.001 followed by Dunn-test with Benjamini-Hochberg p-value adjustment). *L. leonii* exhibited significantly lower values of Tajima’s D, likely reflecting a recent bottleneck in this population. **D)** The distribution of fitness effects (DFE) of new nonsynonymous mutations in *L. perenne* and *L. leonii*. The DFE is visualized in bins of 2*N_e_s* corresponding to the scaled selection coefficient, and the y-axis indicates the proportion of new nonsynonymous mutations expected to fall into each bin. Despite the wide confidence intervals, *L. leonii* has a higher proportion of effectively neutral (−1 < 2*N_e_s* < 1) nonsynonymous mutations and lower proportion of strongly deleterious nonsynonymous mutations (2*N_e_* < - 100) than *L. perenne*.

#### Genomic selfing syndrome in L. leonii?

Population genomic analyses were conducted to test whether loss of distyly in *L. leonii* led to a shift to higher selfing rate, as observed in *L. trigynum* (Gutiérrez-Valencia et al. 2024). All the following statistics were estimated running VCFTools (Danecek et al. 2011) on the filtered VCF files (Suppl. Table S9).

*F_IS_* estimates per population were quite low for both *L. leonii* and *L. perenne* (*L. leonii* mean = 0.19 ± SE = 0.03; *L. perenne* ger2 mean = -0.01 ± SE = 0.02; ger3 mean = -0.02 ± SE = 0.01; ger5 mean = 0.03 ± SE = 0.02) (Figure 4-A), though significantly higher for *L. leonii* compared to all populations of *L. perenne* (Kruskal-Wallis test chi2 = 29.52, p-value < 0.001, Figure 4-A). Assuming equilibrium (Wright 1984), the mean effective self-fertilization rate in *L. leonii* is 0.32 (2 × *F_IS_*/(1 + *F_IS_*)), suggesting that it is mixed-mating (as defined by e.g., Goodwillie et al. 2005). Genome-wide nucleotide diversity (π) was significantly lower for *L. leonii* (mean = 4.05 × 10^-6^ ± SE = 7.3 × 10^-7^) compared to all populations of *L. perenne* (ger2 mean = 4.94 × 10^-4^ ± SE = 9.8 × 10^-6^; ger3 mean = 5.68 × 10^-4^ ± SE = 1.1 × 10^-5^; ger5 mean = 6.42 × 10^-4^ ± SE = 1.2 × 10^-5^) (Kruskal-Wallis test chi2 = 3148.2, p-value < 0.001, Figure 4-B) though differences were significant between all populations (Figure 4-B). Regarding Tajima’s D, we observed the same pattern: significantly lower Tajima’s D estimates for *L. leonii* (mean = -0.10 ± 0.03) compared to all populations of *L. perenne* (ger2 mean = 1.43 ± 0.01; ger3 mean = 1.10 ± 0.01; ger5 mean = 0.90 ± 0.01, Figure 4-C).

To test whether the efficacy of selection differed between the homostylous *L. leonii* and the distylous *L. perenne*, we compared the efficacy of selection on new 0-fold degenerate nonsynonymous mutations (Suppl. Note S2). Despite large confidence intervals due to the limited number of informative segregating sites in the homostylous *L. leonii,* our analyses reveal a reduction in the efficacy of selection, as a significantly higher proportion of new 0-fold degenerate nonsynonymous mutations was inferred to be effectively neutral in *L. leonii* than in *L. perenne* (2*N_e_s* of -1 to 1) (Figure 4-D). We also inferred a lower proportion of highly deleterious new nonsynonymous mutations (2*N_e_s* < -100) in *L. leonii* compared to *L. perenne* (Figure 4-D). A deleterious DFE model was the best fit for *L. leonii*, therefore, beneficial mutations (2*N_e_s* > 1) were only inferred for *L. perenne* (mean proportion of beneficial non-synonymous substitutions DFE_α_ = 0.16).

## DISCUSSION

Loss of distyly has occurred recurrently in flowering plants and is frequently associated with loss of self-incompatibility (SI) and herkogamy (e.g. in *Primula* (Huu et al. 2022) or *Linum* (Gutiérrez-Valencia et al. 2024)), and higher selfing rates (Charlesworth and Charlesworth 1979; Ganders 1979; Barrett 2019; Zhong et al. 2019; Gutiérrez-Valencia et al. 2024). For most of the *Linum* species analysed here, previous studies have shown that distylous species are SI and homostylous species SC (e.g., (Nicholls 1985; Murray 1986; Dulberger 1987; Kearns and Inouye 1994; Gutiérrez-Valencia, P.I. Zervakis, et al. 2024)). The tight association between loss of distyly and SC suggests that this transition might be coupled with a shift to high selfing. According to theory, such shifts would subsequently lead to *N*_*e*_reduction and weaken the efficacy of selection (Charlesworth and Charlesworth 1979; Charlesworth and Wright 2001). However, our results in *Linum* provide mixed evidence for relaxed efficacy of selection in species that have lost distyly, suggesting that a higher selfing rate might not be systematically associated with distyly breakdown.

So far, the main expectation following distyly loss is an increased selfing rate. As previously mentioned, this would ultimately result in relaxation of selection due to reduced *N*_*e*_and reduced effective recombination rate, which would lead to an accumulation of slightly deleterious mutations (Wright et al. 2008; Wright et al. 2013; Cutter 2019). To test whether this was the case in *Linum*, we compared *d*_*N*_⁄*d*_*S*_ values in distylous and homostylous *Linum* lineages. While we found no difference in selective pressure between distylous and homostylous lineages within clade B2, there was a slight relaxation of purifying selection for the homostylous species of clade A, although the difference in *d*_*N*_⁄*d*_*S*_ estimates was quite small and not associated with an increase in *d*_*N*_. The mixed evidence for relaxation of selection in the homostylous species could be the result of the sub-sampling of nuclear genes. Specifically, the angiosperm353 set includes genes chosen to be single-copy and conserved across a wide set of flowering plant species. To the extent that these genes are enriched for strongly constrained sites (which are relatively insensitive to reductions in *N_e_*), our power to detect an effect of selfing might be limited. Yet, working with either 79 (D1) or 129 (D2) genes yielded consistent *d*_*N*_⁄*d*_*S*_ estimates, supporting the robustness of our results. As another explanation, the timing of the shift could affect *d*_*N*_⁄*d*_*S*_ (Wang et al. 2021): increased *d*_*N*_⁄*d*_*S*_ values and signatures of relaxed selective pressure are expected when the shift to higher selfing rate is old, while the opposite pattern is expected with a more recent shift, due to purifying selection acting on the previously heterozygous deleterious mutations (Arunkumar et al. 2015). Thus, the differences in *d*_*N*_⁄*d*_*S*_ estimates between clades might be the result of a difference in timing of the shift to higher selfing rates: species in clade B2 may have transitioned recently to self-compatibility, so that the genomic data would mostly capture the increase in purifying selection rather than the relaxed selective pressures. Additionally, the ASTRAL phylogeny suggests that clade B2 is relatively more recently derived than clade A. While this inference is based on a non-time-calibrated species tree and substitution rate differences, it is consistent with the lower sequence divergence observed in clade B2. The deeper splits observed in the chloroplast tree likely reflect the distinct evolutionary history of the chloroplast genome rather than a true contradiction. Estimating when shifts to selfing occurred using new population genetic methods (e.g., Strütt et al. 2023) and population genomic data for more homostylous species would allow to test that possibility. Finally, as previously mentioned, higher selfing rates might not always accompany the loss of distyly (Yuan et al. 2023), and there is some evidence for additional evolutionary pathways including mixed-mating in *Turnera* (Barrett and Shore 1987), in *Primula* (De Vos et al. 2018), and observed in the field in *Linum arenicola* (Harris and Koptur 2022) and *Linum marginale* (Burdon et al. 1999). In these cases, we would not necessarily expect relaxation of selection. Studies on the rate of outcrossing and the prevalence of SI in additional *Linum* species are needed to test for that.

Chloroplast genome analyses revealed the opposite pattern to that observed in the nuclear genome: no difference between mating systems for clade A species but slightly higher *d*_*N*_⁄*d*_*S*_ values for the homostylous species of clade B2. Similarly, Glémin and Muyle (2014) analysed selective pressure on the chloroplast genes *rbcL* and *matK* in multiple plant groups and found only weak evidence for relaxed purifying selection in the selfing species. The increased *d*_*N*_estimates for homostylous species of clade B2 suggest that this result might be due to moderate relaxation of selection. This would match with a more recent divergence and mating system shift in clade B2 species. In this context, the increase in *d*_*N*_⁄*d*_*S*_ would only be detectable in the chloroplast genome, which is not influenced by dominance effects whereas in the nuclear genomes such a signal would be masked by these, as pointed out in Glémin and Muyle (2014). Additonally, because a genome-wide increase in selfing should similarly affect both genomic compartments, this discrepancy might suggests that the modest nuclear signal observed in clade A might not primarily reflect reduced selection efficiency due to selfing. This supports the view that the loss of distyly can follow multiple evolutionary pathways, not all of which involve substantial increases in selfing and thus, predictable changes in selective pressure.

Consistent with the suggestions of Yuan et al. 2023, our results on the homostylous *L. leonii* provide support for the scenario in which loss of distyly does not systematically lead to increased selfing rates, and demonstrate that evolutionary demographic processes can produce genomic patterns resembling those typically attributed to selfing. We assembled the nuclear genome of *L. leonii* and, integrating population genomic data, tested for genomic effects of loss of distyly through comparison with the distylous close relative *L. perenne*. The two species diverged from each other 1-2 Mya, which sets the approximate upper age limit for loss of distyly in *L. leonii*. This timing broadly corresponds to the major climatic oscillation of the Quaternary (from ∼2.5 Mya to present) which could have played a role in this transition. During this period, dramatic climatic changes led to several shifts in species ranges and speciation events, and could have altered pollinator communities (Davis and Shaw 2001; Kadereit and Abbott 2021). Such changes in pollinator availability and composition may have influenced mating system evolution. In particular, as proposed by Yuan et al. (2023), the transition from long-tongued pollinators to more generalised ones could have a marked effect on distyly and may promote assortative pollen transfer and the evolution of outcrossing or mixed-mating monomorphic individuals. In support of this scenario, *L. leonii* (clade A) which is mixed-mating (selfing rate ∼0.3) has a corolla that is not greatly reduced in size compared to the distylous *L. perenne* (Figure 1). This situation contrasts greatly with that in the selfing homostylous *L. trigynum* (clade B2; selfing rate ∼0.9) which exhibits a floral selfing syndrome with a greatly reduced corolla size compared to its close distylous relative *L. tenue* (Gutierrez-Valencia et al. 2024). The mixed-mating strategy of *L. leonii* contrasts with that of *L. trigynum* and homostylous species in other genera (Zhong et al. 2019; Mora-Carrera et al. 2021; Wang et al. 2021). As even low levels of outcrossing preventprevents accumulation of deleterious mutations (Kamran-Disfani and Agrawal 2014) the mixed-mating *L. leonii* might not be expected to display genomic signatures of selfing.

Nevertheless, similarly to the selfing *L. trigynum* (Gutierrez-Valencia et al. 2024), we observed an overall decrease in genetic diversity and relaxed purifying selection on new nonsynonymous mutations in *L. leonii* compared to *L. perenne.* These results are in agreement with theoretical expectations for relaxed efficiency of selection in populations with reduced *N*_*e*_(Wright 1931; Barton and Partridge 2000; Brandvain and Wright 2016), and resemble the results of mating system transitions in other species (Chen et al. 2017; Laenen et al. 2018; Jaramillo-Correa et al. 2020). Given the lack of evidence for a high selfing rate in *L. leonii*, the reduced *N*_*e*_ and associated genomic signatures are likely to reflect the impact of past demographic processes, such as a prolonged population bottleneck as inferred in our *dadi* analyses, and the geographical location of the sampled populations at the species range margin (Suppl. Fig. S3). *L. leonii* is endemic to France and Germany (Ockendon 1968; McDill et al. 2009; Maguilla et al. 2021) and the sampled population is located at the margin of the current geographical distribution of the species. Populations at the edge of a species’ range may accumulate an expansion load due to allele surfing of deleterious mutations during range expansion (Peischl et al. 2013), this process has already been confirmed in several plant species (González-Martínez et al. 2017; Willi et al. 2018; Zeitler et al. 2023) and could similarly explain the excess of mildly deleterious mutations observed in the sampled population of L. leonii (Suppl. Fig. S3). Range expansion is often thought to confer an advantage to homostylous self-compatible species compared to distylous self-incompatible ones, as their reproductive success does not rely on pollinators, which makes them better colonizers (Baker 1955; Levin 2012). Supporting this idea, a recent study showed an association between range expansion and homostyly in *Linum* (Maguilla et al. 2021). However, this association has not yet been clearly linked to changes in specialised pollinator presence or reductions in outcrossing rates.

## Conclusion

Our study focused on homostylous and distylous species of *Linum* to understand the evolutionary consequences of repeated loss of distyly. Our results show that the evolutionary consequences and genomic signatures associated with distyly breakdown may be more varied than previously thought, as a result of factors such as the timing of the loss, the extent to which homostyly is associated with selfing, and historical population size changes. Our divergence data analyses provide mixed evidence for relaxed selective pressure associated with the loss of distyly and our in-depth study of *L. leonii* demonstrates that loss of distyly is not always associated with a shift to high selfing rates. Instead, we observed genomic signatures of the historical bottlenecks that this species experienced. This result highlights the possibility that a substantial number of homostylous species could instead be mixed-mating, as previously suggested by Yuan et al. (2023), and as such would not be expected to be affected by the long-term negative consequences of selfing, which might ultimately increase the risk of extinction (Takebayashi and Morrell 2001; Igic and Busch 2013). If there is no strong association between loss of distyly and selfing rate across *Linum*, that could explain why no correlation was found between reproductive trait change and extinction rate in this genus (Maguilla et al. 2021). Investigating the varied outcomes of loss of distyly in terms of outcrossing rates and long-term evolutionary and genomic consequences is an important aim for future work.

## MATERIAL AND METHODS

### Testing for relaxed selection in homostylous Linum spp

#### Divergence data generation

To test for relaxed selection in homostylous species of *Linum*, we generated whole-genome sequencing data and target-capture data for sixteen *Linum* species (Suppl. Table S1). We extracted DNA from dried leaf material using the Isolate II Plant DNA Kit (Bioline, UK). Whole-genome sequencing libraries were generated using the TruSeq PCRfree DNA sample preparation kit (cat# 20015962, Illumina) using a target insert size of 350 bp.

For target capture sequencing, library preparation and target capture using the myBaits Angiosperms353 panel (Johnson et al. 2019) was performed by Arbor Biosciences. All sequencing libraries were sequenced on a NovaSeq 6000 to generate paired-end 150 bp reads. We supplemented our sequencing data with publicly available genomic data for three additional *Linum* species and the outgroup species *Tirpitzia sinensis* (Suppl. Table S1). In total, our dataset contained twenty species, among which fourteen were distylous and six were homostylous.

#### Alignment construction

For each sample, chloroplast genomes were independently assembled from whole-genome sequencing data using GetOrganelle v.1.7.3.3 (Jin et al. 2020) with default options and the following k-mer assembly values: 21, 55, 85, 115; and with *--emblant_pt* as the target organelle genome type. Assembly quality was assessed by inspection of organelle genome assembly graphs and running the additional script *summary_get_organelle_output.py* from the GetOrganelle pipeline. Chloroplast genome annotation was conducted online using the GeSeq v2.03 software for rapid annotation of organellar plant sequences (Tillich et al. 2017). GeSeq conducts standard BLAT analyses against a high quality manually curated reference database spanning a wide range of plant taxa to fully annotate genomes (i.e. protein coding genes, introns and exons, as well as transfer RNA (tRNA) and ribosomal RNA (rRNA)). Visual chloroplast genome maps were done using OGDRAW (Greiner et al. 2019). We retrieved 112/115 chloroplast genes from the reference database of GeSeq. Based on the GeSeq annotation, we further checked the identified inverted repeat (IR) for each assembly by comparing CDS sequences of genes located in the IR using MASH distance estimation v2.3 (Ondov et al. 2016). Sequences of such genes should be 100% identical and mash distance should hence be close to 0. We then specifically extracted CDS sequences for each of the 112 annotated genes and constructed gene alignments only for those containing sequences for each species (i.e. 106/112 genes) using SeqKit v0.15.0 (Shen et al. 2016) and codon-based alignment T-Coffee v11.0 (Notredame et al. 2000). We kept only the 101 genes for which alignments were of good quality (i.e., T-Coffee alignment score > 80 and no internal stop codons). We further removed the *trn* and *rrn* genes so the rest of the analyses were conducted on the remaining 67 chloroplast genes.

For the nuclear data, we used Hybpiper v2.1.6 (Johnson et al. 2016) to obtain per gene fasta files. Briefly, Hybpiper is a python pipeline designed to construct gene files from sequence capture data using a target file containing gene sequences of genes of interest. It first mapped the reads to the reference targeted genes using BLASTx (Camacho et al. 2009), DIAMOND (Buchfink et al. 2015) or BWA (Li and Durbin 2009). Then, the mapped reads were assembled specifically for each gene of interest using SPAdes (Bankevich et al. 2012). The final output was a fasta file which included sequences for each gene of interest. We ran Hybpiper with the recommended options (i.e., *--run_intronerate --timeout_assemble* 200 *--cpu* 20 *--no_padding_supercontigs --single_cell_assembly*) using the target file provided for the 353-probe dataset (Johnson *et al*. 2019). Hybpiper output were further analysed using the “*hybpiper stats*”, “*hybpiper recovery_heatmap*” and “*hybpiper retrieve_sequences*” scripts from Hybpiper GitHub (https://github.com/mossmatters/HybPiper). We used the target file associated with the 353 DNA dataset (Angiosperm353.fasta, https://datadryad.org/stash/dataset/doi:10.5061/dryad.s3h9r6j). The sequences of *T. sinensis, L. bienne, L. decumbens* and *L. lewisii* were already in the 353 datasets, from the OneKP project (Wickett et al. 2014) so we did not run Hybpiper for these species and instead retrieved the corresponding coding sequences from NCBI (Suppl. Table S1). Not all genes were retrieved for all species, and some genes were also identified as containing potential paralogs (Suppl. Table S12). To reduce complications from missing data, we only analysed the 142/353 nuclear genes that contained gene sequences for all twenty species. We constructed codon-aware alignments for these 142 genes using SeqKit and T-coffee. Out of these genes, we further selected only the 129/142 with high-quality alignments, following the same criteria as for the chloroplast data set. To control for the paralog warning from Hybpiper without losing too many genes, we further divided the nuclear data into two gene sets: “D1” containing the 79/129 genes without any paralog warning from hybpiper and the “D2” dataset, containing all 129 genes, including those with a paralog warning. All downstream analyses were run on both the D1 and D2 gene sets.

#### Phylogeny reconstruction

Before estimating selective pressure on the chloroplast and nuclear datasets, we reconstructed a chloroplast-based phylogeny. To do so, we concatenated the 67 identified chloroplast genes, and reconstructed phylogenetic trees using RAxML v8.2.12 (Stamatakis 2014) and a GTR GAMMA model of substitution. We additionally specified the outgroup (*Tirpitzia sinensis*) using the option *-otg*. We performed 1000 replicates of rapid bootstrap analysis (option *-f a*), followed by a search for the best-scoring maximum-likelihood (ML) tree *(-# 1000)*. Each bootstrap replicate involved a single ML search. The chloroplast tree was further used for analyses of selection on the chloroplast genes and for ancestral state reconstruction.

To test whether the topology of the chloroplast tree was also recovered for the nuclear dataset, we first built gene trees for all genes from the D2 dataset as above for the chloroplast dataset. We then used these individual gene trees to estimate a single species tree using ASTRAL v.5.7.8 (Zhang et al. 2018). We additionally ran RAxML with the same options as for the chloroplast dataset on the concatenation of the D2 dataset to get node support and branch length for the ASTRAL phylogeny, as suggested in the ASTRAL manual. We reported the results only of the ASTRAL phylogeny built with the D2 dataset. The ASTRAL topology was used for analyses of selection on nuclear genes.

#### Ancestral state reconstruction

To be able to perform ancestral state reconstruction, we time-calibrated the chloroplast phylogenetic tree, using the R package ape v5.8 (Paradis and Schliep 2019). Chronograms were generated using the chronos() function and a correlated rate model and calibrating the root node based on the age of the most recent common ancestor of *Linum* and *T. sinensis*, following Maguilla et *al*. (2021). To account for unobserved evolutionary processes potentially influencing the underlying evolution of different state histories, we reconstructed ancestral states using the Hidden rates model (Beaulieu et al. 2013), which incorporates hidden states in the analyses, as implemented in the R package corHMM v2.8 (Boyko and Beaulieu 2021). Initially we ran corHMM and fitted separate models with one, two, and three rate categories. However, models incorporating multiple rate categories showed no significant improvement based on the Akaike Information Criterion (AIC). Specifically, the rate of transitioning from homostyly to distyly (reversal to distyly in our case, which is equivalent to the gain of a complex character) was estimated at a rate three times higher than the forward transition (distyly to homostyly) under the simplest, best-supported model. As shifts from SI to SC generally occur much more frequently than the reverse transition (Igic et al. 2006; Wright et al. 2013), we considered these rate estimates to be biologically implausible. We therefore instead defined new transition rates matrices and repeated our analyses with corHMM: transition rates from distyly to homostyly varied between 1 × 10^-10^ and 1 in 10-fold increments, while the rates for the opposite transition were set to be consistently lower, from 1 ×10^-100^ to 1 × 10^-4^. In all cases, we fixed the root of the tree (the *Linum* – *Tirpitzia* common ancestor) and the root of *Linum* to distyly to reflect shared ancestral polymorphism at the *S-*locus of *Linum* dating back to the origin of the genus (Zervakis, Postel et al. 2025). After finding the best model based on AICc, we used phytools in R (Revell 2024) to infer the ancestral state of each node using 1000 reconstructions with makeSimmap(). The final results were visualized with plotSimmap().

#### Divergence-based analyses of selection

To test for an effect of loss of distyly on selection, we separately estimated *d*_*N*_⁄*d*_*S*_ (a proxy for selective pressure) for chloroplast and nuclear gene alignments using PAML – codeml v4.9 (Yang 2007). We ran codeml using the Branch-Model, allowing the *d*_*N*_⁄*d*_*S*_ to vary along specific annotated branches and then comparing the null model (i.e. H0, where the whole tree shared the same *d*_*N*_⁄*d*_*S*_) with different branch-model annotated trees (Suppl. Table S3). Annotation of internal branches of the phylogenetic tree as distylous or homostylous was based on the ancestral state reconstruction. To select the best supported hypothesis and estimation of *d*_*N*_⁄*d*_*S*_, we used Log-Likelihood Ratio Tests (LRT) as suggested in (Yang 2007).

For the chloroplast dataset, we ran PAML on the set of 67 concatenated chloroplast gene alignments that was also used to construct the chloroplast phylogenetic tree. For the nuclear dataset, we ran codeml on both D1 and D2 datasets. As results were highly similar for the two datasets, we will only report results of D2, the larger dataset, in the main text. Results for D1 can be found in supplementary table S4 and S5. For both D1 and D2, we used the unrooted ASTRAL nuclear tree topology.

We additionally extracted the separate estimates of *d*_*N*_ and *d*_*S*_ for each tip branch from the best branch-model, to further decipher whether differences in *d*_*N*_⁄*d*_*S*_ could be the result of higher *d*_*N*_ and changes in selective pressure. We then computed means per clade and within clade, per mating system. To test for significant differences of *d*_*N*_ and *d*_*S*_ between these groups, we conducted a Kruskal-Wallis on R. The estimates are reported in table 3.

### Characterizing the evolutionary consequences of loss of distyly in L. leonii

#### Genome sequencing, assembly and annotation

High-molecular-weight (HMW) DNA from one *L. leonii* individual (LEO-2-29-2; Figure 1-A) was used to prepare SMRTbell libraries for high-fidelity (HiFi) long-read sequencing. Each library was run on two SMRT cells in HiFi mode using the Sequel II system (Pacific Biosciences), yielding 39.5 Gb of HiFi data with an average read length of 15 kbp. To produce high-quality proximity ligation (Hi-C) libraries for genome assembly scaffolding, 300 mg of fresh-frozen leaf tissue was ground to a fine powder for preparation of Hi-C libraries using the Dovetail Omni-C Kit. Sequencing on Illumina NovaSeq 6000 platform generated approximately 0.97 × 10⁹ paired-end 150 bp reads.

For genome annotation, RNA sequencing data were collected from leaves, stems, flower buds, and open flowers of *L. leonii* LEO-2-29-2. Total RNA was extracted using the RNeasy Plant Mini Kit (Qiagen). Sequencing libraries were prepared with the TruSeq Stranded mRNA Library Preparation Kit (Illumina, San Diego, CA, USA), including poly(A) selection and unique dual indexes. Libraries were sequenced using paired-end 150 bp reads on the NovaSeq 6000 platform.

We generated primary and haplotype-resolved genome assemblies for the 2-29-2 individual using HiFi and Hi-C data, employing the integrated Hi-C assembly mode in Hifiasm (Cheng et al. 2021). The primary assembly was used for all downstream analyses (Suppl. Table S6). Assembly completeness was assessed using Benchmarking Universal Single-Copy Orthologs (BUSCO; Waterhouse et al. 2018) with the *eudicots_odb10* gene set. Prior to annotation, assemblies were screened for contamination and for chloroplast and mitochondrial sequences, following the approach described by Gutiérrez-Valencia et al. (2024). Four regions (∼ 834 kbp) were hard masked from the primary assembly (using “N”s) due to signs of contamination: ptg000010l: 25960200-26058400, ptg000012l: 1528800-1774000 and 5840800-5929200, ptg000013l: 30311400-30713400. Genome annotation was performed by the National Bioinformatics Infrastructure Sweden (NBIS) Annotation and Assembly unit as in Zervakis and Postel et al. (2025). Transposable element annotation was performed using HiTE v3.2 (Hu et al. 2024).

#### Whole-genome sequence data from natural populations

We generated whole-genome short-read sequencing data from 20 individuals of *L. leonii* from a natural population in Germany (Suppl. Table S8) as described above in the “*Divergence Sequencing Data*”. To contrast population genetic patterns in the homostylous *L. leonii* to those in a closely related distylous species, we used whole-genome sequences from 136 individuals from three natural populations in Germany of the distylous *L. perenne* (Zervakis, Postel et al. 2025, Suppl. Table S8). We further used Illumina whole-genome resequencing reads from one individual of *L. grandiflorum* (Lgra-62-06) to retrieve outgroup information (Zervakis, Postel et al. 2025).

#### Bioinformatic processing of population genomic data

We adapter-trimmed Illumina whole-genome sequencing reads using BBDuk from BBMap v38.61b (Bushnell 2014). *L. leonii* reads were mapped to the primary genome assembly of *L. leonii* and *L. perenne* reads to our published *L. perenne* assembly (Zervakis, Postel et al. 2025), except for analyses of population structure and demographic history, for which all reads were mapped to the *L. perenne* assembly only (Suppl. Table S9). We used BWA-MEM v.0.7.17 (Li 2013) for read mapping, including only proper pairs (FLAG 0×2) and excluding reads with mapping quality (MAPQ) lower than 20. We removed duplicate reads with Picard MarkDuplicates v2.20.4 (http://broadinstitute.github.io/picard). We called variants using BCFTOOLS mpileup v.1.17. (Danecek et al. 2021). Variants were filtered in the following way, similarly to Zervakis and Postel et al. (2025): (i) only bi-allelic variants and invariant sites were kept, (ii) we applied filters for depth, missingness and mapping quality (BCFtools min_depth = 5; max_depth = 200; missingness = 0.9; min_quality = 20), (iii) additional masking of repeats was necessary due to the high repeat content of our assemblies so we masked repeats using ‘*bedtools intersect*’ and by filtering on coverage as in Gutiérrez-Valencia et al. (2021) and (vi) we applied an allele balance filter with thresholds 0.2 and 0.8, setting heterozygous calls that failed this criterion to missing, in order to reduce false heterozygous calls.

#### Population genomic analyses of inbreeding and polymorphism

To assess whether the loss of distyly in *L. leonii* was associated with a higher selfing rate and a lower nucleotide diversity, we ran the following analyses. We first estimated the inbreeding coefficient *F_IS_*(Wright 1949), nucleotide diversity (*π*), and Tajima’s D genome-wide and in window sizes of 100 kb in VCFTools (Danecek et al. 2011). We tested for a difference in *F_IS_*, π and Tajima’s D between populations and species using a Kruskal-Wallis test followed by post-hoc Dunn-test with Benjamini-Hochberg p-value adjustment in R (R Core Team 2021).

#### Population structure and demographic history inference

To test for population structure and gene flow in *L. perenne* and *L. leonii*, we ran a Principal Component Analysis (PCA) as well as an ADMIXTURE analysis (Alexander et al. 2009). We used PLINK (Chang et al. 2015) to prune the VCF based on linkage disequilibrium (*LD*) using the pairwise squared correlation coefficient (*r*^2^) with the option *--indep-pairwise* 50kb 1 0.2 (window size in kb= 50, variant count to shift the window at the end of each step = 1, pairwise *r*^2^ threshold = 0.5), and subsequently ran the PCA using the *-pca* option. ADMIXTURE analysis (Alexander et al. 2009) was conducted on the *LD*-pruned dataset with K values ranging from 1 to 9 and using the *-CV* (Cross-Validation) option to choose the best K value (i.e. the one with the lowest cross validation value). Both PCA and ADMIXTURE results were plotted in R. To investigate the demographic history of each population, we used *dadi* models with inbreeding (Gutenkunst et al. 2009; Blischak et al. 2020), a diffusion approximation-based method that estimates demographic parameters and inbreeding coefficients by maximizing the likelihood from site frequency spectra (SFS). We first ran fastdfe v1.1.9 (Sendrowski and Bataillon 2024) to infer the ancestral state of each allele and polarise the SFS, using *L. grandiflorum* (Zervakis, Postel et al. 2025) as an outgroup. Specifically, we used the class “MaximumLikelihoodAncestralAnnotation”, which annotates the vcf using a maximum likelihood approach based on the probabilistic model of est-sfs (Keightley and Jackson 2018). We then annotated 0- and 4-fold sites in the *L. leonii* and *L. perenne* genomes using the “degenotate.py” python script (https://github.com/harvardinformatics/degenotate).

Subsequently, we selected the 4-fold sites and we generated unfolded single and joint population SFS from high-quality biallelic SNPs using easySFS (https://github.com/isaacovercast/easySFS), employing the “*-a*” option to retain all variant and invariant sites. Invariant sites were included to scale the *dadi* output (*N_e_anc_* = *θ* / (4 * *μ* * *L*), where *θ* is nucleotide diversity as inferred by *dadi*, *μ* is the assumed mutation rate, and *L* is the sum of invariant and variant sites). For each analysis, allele counts were down sampled (Suppl. Table S11) to maximise the number of variant sites and mitigate the effects of missing data in populations with low nucleotide diversity. Dadi was ran only using 4-fold sites.

We applied *dadi* to joint SFSs in two-population (2D) models, for *L. leonii* vs each of *L. perenne* populations, to reconstruct the timing and parameters related to the species split. Based on our ADMIXTURE results, we assumed population splits without gene flow, followed by scenarios involving stable or varying population sizes (six scenarios in total for 2D models; Suppl. Figure S5). Each model was optimized through at least 50 rounds of 300 iterations using the *dadi.Inference.optimize_log_fmin* optimizer, and we used AIC to select the models with the best fit. To scale generation times into years we assumed a generation time of 2 years in *L. perenne*, based on the long lifespan, yet limited seed set of *L. perenne* during the first year (Ockendon 1968; Tork et al. 2024) and the ancestral reconstruction results suggesting an outcrossing ancestor. We assumed a generation time of 1 year in *L. leonii* due to its shorter lifespan (Ockendon 1968). Finally, we assumed a mutation rate of 7.1 × 10^-9^ based on sequencing of mutation accumulation lines of *Arabidopsis thaliana* (Ossowski et al. 2010). Results are detailed in the manuscript for *L. leonii* vs *L. perenne* - ger2. Because results were similar between all three comparisons, the outputs for the two other pairs can be found in Suppl. Note S1.

#### Population genomic analyses of selection

To test whether loss of distyly in *L. leonii* had an impact on the efficacy of selection, we estimated the distribution of fitness effects (DFE) of new nonsynonymous (0-fold degenerate) mutations. We analysed unfolded SFS for 0-fold and 4-fold sites in fastdfe v1.1.9 (Sendrowski and Bataillon 2024) to estimate the DFE of new 0-fold degenerate nonsynonymous mutations.

First, we tested all the provided parameterizations and checked nested models for presence of beneficial mutations and ancestral variant misidentification using the Akaike Information Criterion (AIC). After choosing the best model for each population, we inferred the DFE and the proportion of new beneficial 0-fold degenerate non-synonymous substitutions (*α*) using 100 iterations and 1000 bootstraps. Finally, we rescaled the discretized results to 2*N_e_s* instead of the default 4*N_e_s* and visualized our results in python.

## Data availability

All sequencing data generated in this study will be made publicly available via the European Nucleotide Archive (ENA) upon publication.

## Acknowledgments

We thank Mohamed Abdelaziz, Benjamin Laenen and Aurélie Désamoré for help with sampling. This work was supported by funding from the European Research Council (ERC) under the European Union’s Horizon 2020 and Horizon Europe research and innovation programs [grant agreement nos.: 757451 and 101132305], from the Swedish Research Council [grant agreement nos.: 2019-04452 and 2023-04532] and from the Erik Philip-Sörensen foundation to [TS] and the Nilsson-Ehle foundation to [PIZ]. [ZP] was funded by a Carl Tryggers foundation [grant CTS21:1471] and by a grant from the Sven and Lily Lawski foundation. Computations were enabled by resources provided by the National Academic Infrastructure for Supercomputing in Sweden (NAISS), partially funded by the Swedish Research Council [grant agreement no. 2022-06725]. We thank the PDC Center for High Performance Computing (KTH Royal Institute of Technology) for providing the computing resources (Dardel) and related services that have been used in this work. Short-read library construction and sequencing was performed by the SNP&SEQ Technology Platform in Uppsala. The facility is part of the National Genomics Infrastructure (NGI) Sweden and Science for Life Laboratory. The SNP&SEQ Platform is also supported by the Swedish Research Council. The authors would like to acknowledge support of the National Genomics Infrastructure (NGI) / Uppsala Genome Center and UPPMAX for providing assistance in massive parallel sequencing and computational infrastructure. Work performed at NGI / Uppsala Genome Center has been funded by RFI / VR and Science for Life Laboratory, Sweden. This work was supported by the SciLifeLab & Wallenberg Data Driven Life Science Program, Knut and Alice Wallenberg Foundation (grants: KAW 2020.0239 and KAW 2017.0003), and by the National Bioinformatics Infrastructure Sweden (NBIS) at SciLifeLab.

