## Supplementary material for "Evolutionary consequences of repeated loss of distyly in *Linum*"

#### Supplementary Figures


**Figure S1 –** Example of a reconstructed chloroplast genome assembly

**Figure S2 –** ASTRAL phylogeny based on the 217 gene trees

**Figure S3** – Geographical distribution of *Linum leonii* and *Linum perenne*

**Figure S4** – PCA plots with both species' populations

**Figure S5** – Demographic history 2D models


#### Supplementary tables


**Supplementary table S1** – Sample information per species

**Supplementary table S2** – GetOrganelle and GeSeq annotation metrics for the chloroplast genome assembly of each species

**Supplementary table S3** – Hypothesis tested with PAML-codeml, Branch-model

**Supplementary table S4** – Log likelihood-ratio test (LRT) results for D1 nuclear genes

**Supplementary table S5** – $d_N/d_S$ values for D1 nuclear genes, estimated using PAML-codeml, Branch-model H2

**Supplementary table S6** – *Linum leonii* genome assembly metrics

**Supplementary table S7** – *Linum leonii* genome assembly transposable elements (TEs) composition.

**Supplementary table S8** – Sample information for *Linum leonii* and *Linum perenne* population genomic analyses

**Supplementary table S9** – Read mapping information

**Supplementary table S10** – ADMIXTURE cross-validation values

**Supplementary table S11** – *dadi* 2D models analysis parameters

**Supplementary table S12** – Hybpiper statistics


#### Supplementary dataset


**Supplementary dataset S1** – Genome assembly report for *Linum leonii*


#### Supplementary Notes

**Supplementary Note S1 -** Demographic history inference for 2D SFS between *L. leonii* and populations of *L. perenne* ger3 and ger5.

**Supplementary Note S2** – fastDFE best supported models

### SUPPLEMENTARY FIGURES

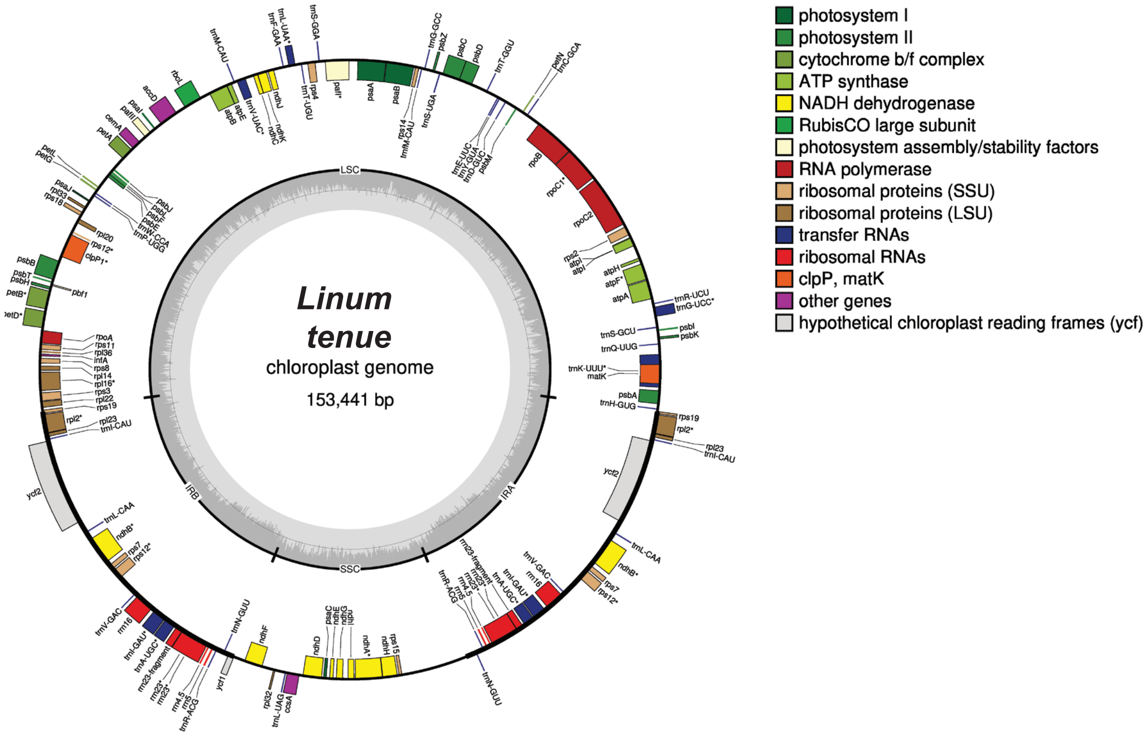


**Figure S1 – Example of a reconstructed chloroplast genome assembly.** The GC content is displayed on first inner circle. The limits of the Small and Large Single Copy Region (SSC and LSC) as well as limits of the inverted repeat (IR) can be found on the second inner circle. Gene's length and position can be retrieved on the chloroplast genome maps third circle. Each gene is colored depending on its chloroplast complex. The chloroplast genome size is indicated at the center of the map. Sizes of the different parts of the chloroplast genome (i.e. LSC, SSC, IR) can be found in Suppl. Table S2.

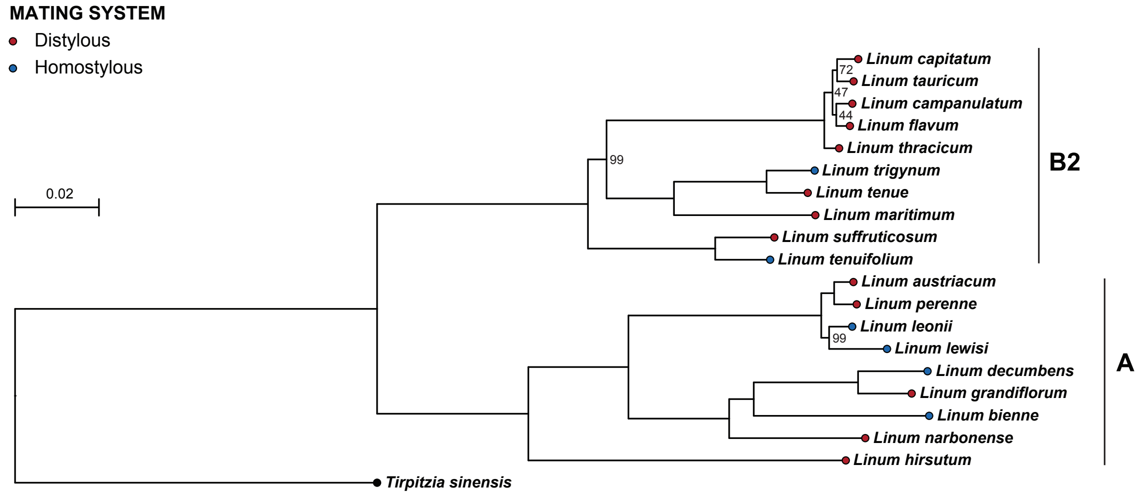


**Figure S2 - ASTRAL phylogeny based on the 129 gene trees.** This ASTRAL phylogeny was built using the 129 gene tree obtained with RAxML. Branch length and node were obtained running RAxML with a GTR GAMMA model of nucleotide substitution and a rapid bootstrap analysis with 1000 replicates on the tree built with the concatenation of the 129 genes (i.e. D2 dataset). Only nodes support with bootstrap value <100 are displayed. Species phylogenetic relationships are the same than the one observed with the chloroplast phylogeny. The only difference is the positioning of L. thracicum in clade B2 but the node support is quite low suggesting that this node is not well resolved as for the chloroplast data.

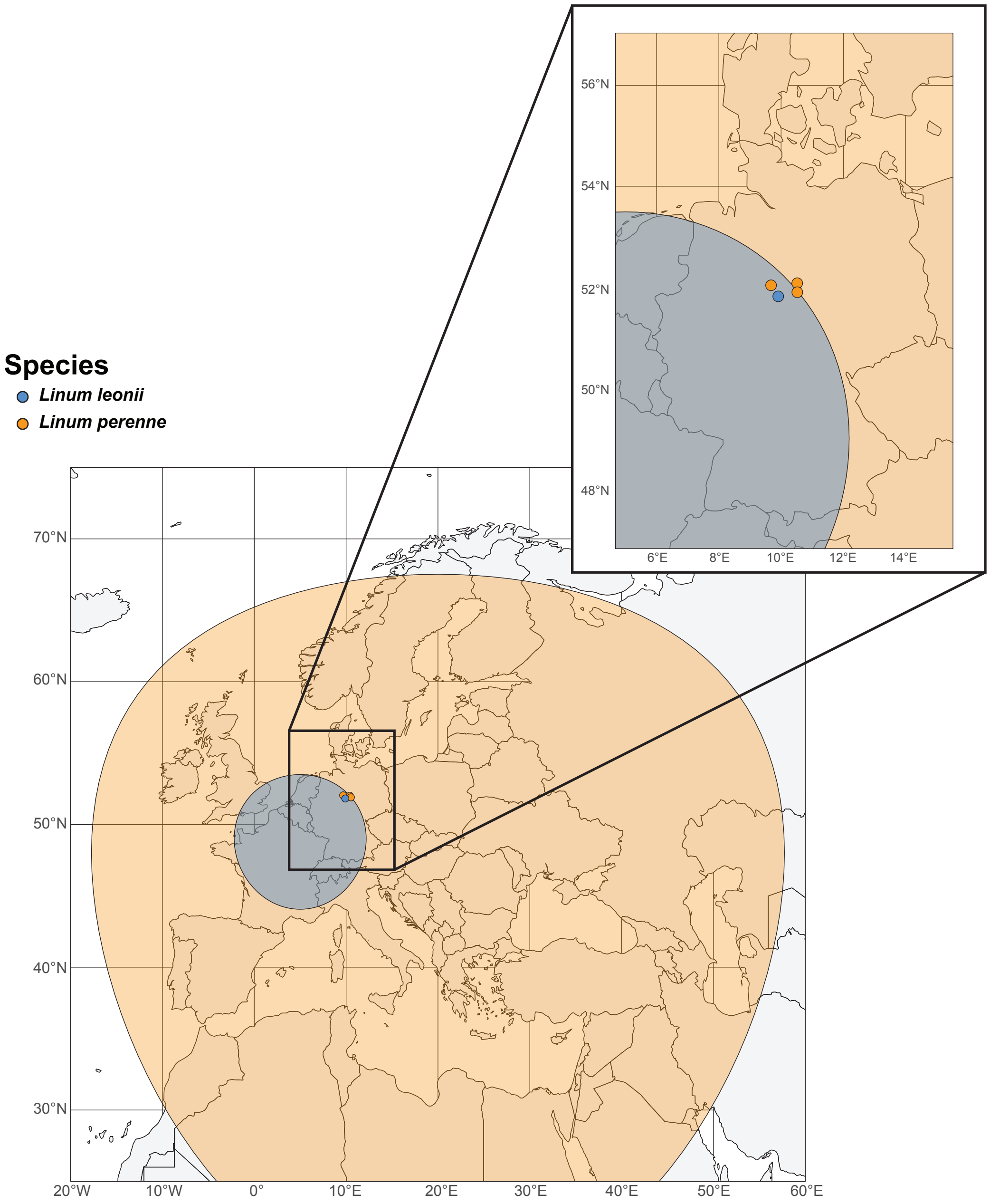


**Figure S3 – Geographical distribution of *Linum leonii* and *Linum perenne*.** *L. leonii* is represented in blue and *L. perenne* in orange. Sampled populations for both species are displayed and their geographical location zoomed in in the upper panel. Data were collected from GBIF and refined based on previous publications (Ockendon 1968, McDill et al. 2009, Maguilla et al. 2021). The figure was built on R.

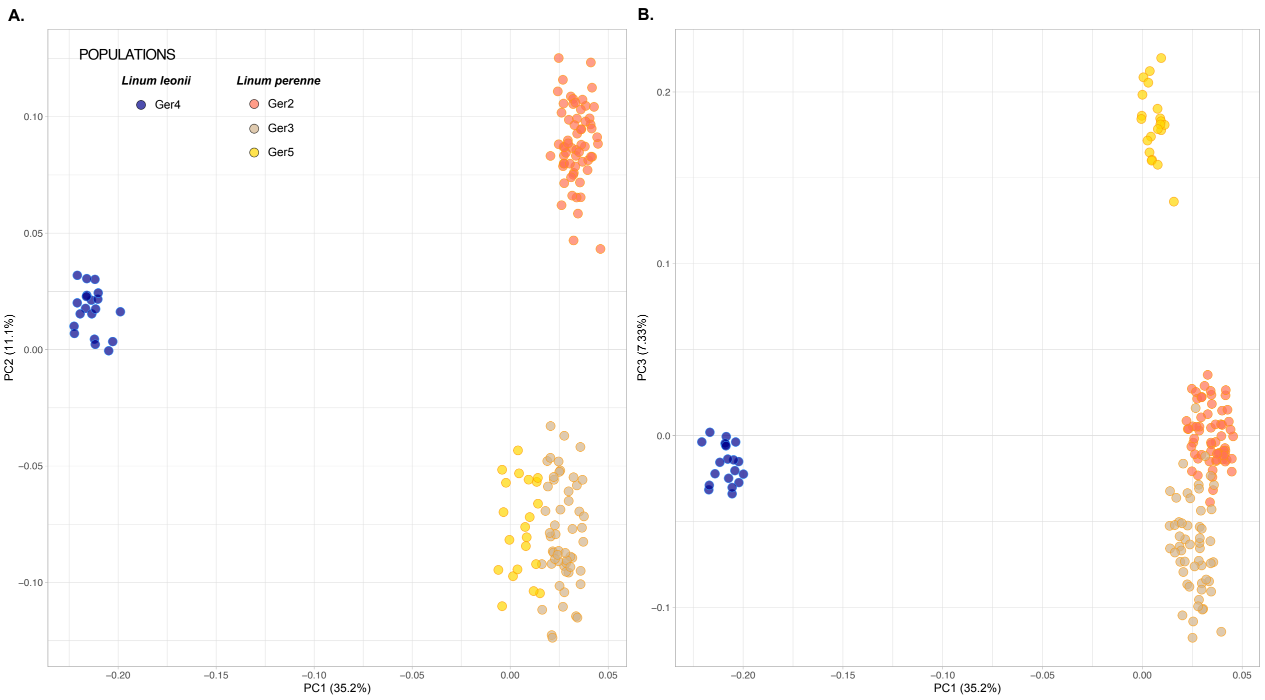


**Figure S4 – Principal Component Analysis (PCA) plot with both species' populations: (A) PC1 and PC2; (B) PC1 and PC3.** Individuals correctly clustered within each population. Additionally, both species separated along PC1 and PC2. *L. perenne*'s population ger2 differentiate from the other ones along PC2 and population ger5 differentiate along PC3 suggesting that population ger3 might be slightly admixed with both ger2 and ger5.

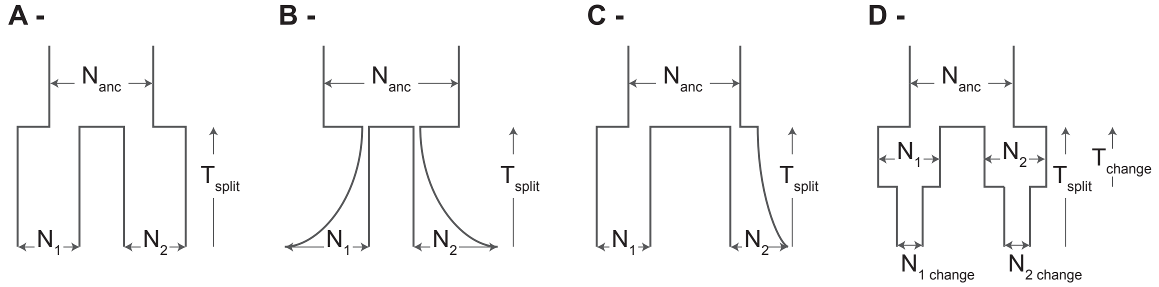


**Figure S5 - Demographic history 2D models.** Schematic representation of the tested demographic history models for joint site frequency spectra. The exponential models (C, D) can represent gradual increase or decrease in effective population size. Abbreviations are as described in Suppl. Table S11. The models' descriptions are the following:

- *split_no_growth* **(A)**: Split at T_split; Immediately two constant-size populations (N1, N2) without growth; Inbreeding ($F_1$, $F_2$).
- *vic_nomig* **(A)**: Split at T_split with proportions s and (1-s); Immediately two constant-size populations (N1, N2) without growth; Inbreeding ($F_1$, $F_2$).
- *split_growth* **(B)**: Split at T_split; Two exponential growing populations; Inbreeding ($F_1$, $F_2$).
- *split_growth_s* **(B)**: Split at T_split with proportions s and (1-s); Two exponentially growing populations; Inbreeding ($F_1$, $F_2$).
- *Founder_nomig* **(C)**: Split at T_split; pop2 starts at fraction s of ancestral Ne, pop1 at 1–s; Pop2 grows exponentially to N2; pop1 stays constant at N1; Inbreeding ($F_1$, $F_2$).
- *no_mig_size* **(D)**: Split at T_split; after split, each daughter population has constant size (N1, N2); Then at time T_change, both change to sizes (N1_change, N2_change); Inbreeding ($F_1$, $F_2$).

### SUPPLEMENTARY TABLES

#### *Legends*

**Supplementary table S1 – Sample's information per species.** Samples' information for all *Linum* species used to estimates selective pressure in the homostylous vs distylous species. This tables report all information such as the source of the sample, the NCBI reference for chloroplast and nuclear data for the species that were already publicly available, the taxonomic section of the sample, its mating system and life style.

**Supplementary table S2 – GetOrganelle and GeSeq annotation metrics for the chloroplast genome assembly of each species.** Plastome assemblies' metrics and annotation for all twenty species. This table reports k-mer and assembly coverage as well as the number of mapped reads and whether GetOrganelle managed to retrieve the circular structure of the plastome assembly. We are also reporting the size of the plastome assembly as well as the size of its different parts (long and small single copy and the inverted repeat).

**Supplementary table S3 – Hypothesis tested with PAML-codeml, Branch-model.** Explanation of the different nested model ran on PAML – codeml. H0 correspond to the unannotated phylogenetic tree, thus with only one estimation of the $d_N/d_S$ ratio for the whole tree. H1 correspond to the phylogenetic tree annotated with three different groups: distylous / homostylous / background, so with three independent estimations of the $d_N/d_S$ ratio (i.e. one per group). H2 corresponds to the annotation of the phylogenetic tree into five different groups: clade A – distylous / clade A – homostylous / clade B2 – distylous / clade B2 - homostylous / background, thus with five different estimations of the $d_N/d_S$ ratio.

**Supplementary table S4 – Log likelihood-ratio test (LRT) results for D1 nuclear genes.** Results of the log-likelihood ratio test for the nuclear D1 dataset. The LRT were done between H0 vs H1 and then H1 vs H2. As for the D2 dataset, the best branch model was H2 (i.e. one $d_N/d_S$ ratio estimated per clade and per mating-system).

**Supplementary table S5 – $d_N/d_S$ values for D1 nuclear genes, estimated using PAML-codeml, Branch-model H2**. Estimated $d_N/d_S$ ratio based on the H2 branch model, for the D1 nuclear genes set. As for the D2 dataset, clade A homostylous species exhibit slightly elevated $d_N/d_S$ ratio compared to the distylous species while no differences are observed between homostylous and distylous species of clade B2.

**Supplementary table S6 – *Linum leonii* genome assembly statistics.** Report of the main metrics for the newly built genome assembly of *L. leonii*. The assembly of length 889 Mb is of good quality, with a BUSCO score of 93.9%.

**Supplementary table S7 – *Linum leonii* genome assembly transposable elements (TEs) composition.**

**Supplementary table S8 – Samples information for *Linum leonii* and *Linum perenne* population's genomic analyses**. This table reports information of the sampling location of the samples used to test for the genomic selfing in syndrome in the homostylous *L. leonii* in contrast to its distylous relatives, *L. perenne*. It also reports the number of samples for each population.

**Supplementary table S9 – Read mapping information to get the different VCF files.** This tables details which reads were mapped to which assembly for each analysis. The VCF files were then built based on these mapped reads.

**Supplementary table S10 – ADMIXTURE cross-validation values.** Reports of the different CV values for each cluster number (K). The lowest CV value was retrieved for K8 indicating that *L. leonii* and *L. perenne* individuals' group into eight different genetic cluster.

**Supplementary table S11 – *dadi* 2D best-models' parameters.** Report of the different parameters for dadi 2D best-fitting models for comparison between *L. leonii* and each population of *L. perenne*. Schematic representations of the models are shown in Suppl. Fig. S5.

**Supplementary table S12 – Hybpiper statistics**. This table reports the main statistics from Hybpiper pipeline, such as the total number of reads and the number of mapped reads and the number of genes with a paralog warning, per species.

#### Suppl. Table S1 – Samples' information per species.

| Species | Source | Chloroplast data | Nuclear data | Section* | Mating system | Clade | Life style** |
|---|---|---|---|---|---|---|---|
| *Linum hirsutum* | LIN 2100 (IPK) | Short read, WGS | Target Capture | Dasylinum | Distylous | Clade A | Perennial |
| *Linum austriacum* | LIN 1883 (IPK) | Short read, WGS | Target Capture | Linum | Distylous | Clade A | Perennial |
| *Linum grandiflorum* | LIN 974 (IPK) | Short read, WGS | Short read, WGS | Linum | Distylous | Clade A | Annual |
| *Linum narbonense* | LIN 1807 (IPK) | Short read, WGS | Target Capture | Linum | Distylous | Clade A | Perennial |
| *Linum perenne* | LIN 2003 (IPK) | Short read, WGS | Target Capture | Linum | Distylous | Clade A | Perennial |
| *Linum bienne* | - | SRR1592607 | SRR1592607 | Linum | Homostylous | Clade A | Annual |
| *Linum decumbens* | - | MV366795.1 | SRR1592610 | Linum | Homostylous | Clade A | Annual |
| *Linum leonii* | 18-2, ger2, 2018 near Moringen, Germany | Short read, WGS | Target Capture | Linum | Homostylous | Clade A | Perennial |
| *Linum lewisii* | - | NC_058799.1 | SRR1592654 | Linum | Homostylous | Clade A | Perennial |
| *Linum thracicum* | LIN 1553 (IPK) | Short read, WGS | Target Capture | - | Distylous | Clade B2 | Perennial |
| *Linum maritimum* | MAR-12, Soler botanical garden, Spain | Short read, WGS | Target Capture | Linopsis B | Distylous | Clade B2 | Perennial |
| *Linum tenue* | Lten-21-22, SAMEA114529652 | Short read, WGS | Target Capture | Linopsis B | Distylous | Clade B2 | Annual |
| *Linum trigynum* | Ltri-10-7, SAMEA114529727 | Short read, WGS | Target Capture | Linopsis B | Homostylous | Clade B2 | Annual |
| *Linum suffruticosum* | 18-5-2 near Ronda, Spain | Short read, WGS | Target Capture | Linopsis C | Distylous | Clade B2 | Perennial |
| *Linum tenuifolium* | LIN 1759 (IPK) | Short read, WGS | Target Capture | Linopsis C | Homostylous | Clade B2 | Perennial |
| *Linum campanulatum* | LIN 1760 (IPK) | Short read, WGS | Target Capture | Syllinum | Distylous | Clade B2 | Perennial |
| *Linum capitatum* | LIN 1903 (IPK) | Short read, WGS | Target Capture | Syllinum | Distylous | Clade B2 | Perennial |
| *Linum flavum* | LIN 2001 (IPK) | Short read, WGS | Target Capture | Syllinum | Distylous | Clade B2 | Perennial |
| *Linum tauricum* | LIN 1658 (IPK) | Short read, WGS | Target Capture | Syllinum | Distylous | Clade B2 | Perennial |
| *Tirpitzia sinensis* | - | ON881454.1 | ERR9230207 | - | Distylous | Outgroup | Na |

*Maguilla et al. 2021
**References for mating system and life habits: Ruiz-Martin et al. 2018, Maguilla et al. 2021, Valdes-Florido et al. 2023, Tork et al. 2021, Jahnke and Etterson. 2018, Link

#### Suppl. Table S2 – GetOrganelle and GeSeq annotation metrics for the chloroplast genome assembly of each species.

| **Clade** | **Mating system** | **Species** | **Kmer cov** | **Asm cov** | **Nb mapped reads** | **Graph (K115)** | **Size – in bp** | | | |
|---|---|---|---|---|---|---|---|---|---|---|
| | | | | | | | **Total asm** | **LSC** | **IR** | **SSC** |
| Clade A | Distylous | *L. austricanum* | 116.4 | 475.0x | 271,381 | Circular | 172,330 | 72,208 | 44,567 | 10,988 |
| | | *L. grandiflorum* | 130.7 | 533.3x | 283,580 | Circular | 158,174 | 82,760 | 32,480 | 10,454 |
| | | *L. hirsutum* | 123.5 | 504.1x | 276,026 | Circular | 165,215 | 79,881 | 38,237 | 8,860 |
| | | *L. narbonense* | 124.3 | 507.2x | 272,110 | Circular | 159,233 | 83,040 | 32,665 | 10,863 |
| | | *L. perenne* | 117.2 | 478.1x | 273,028 | Circular | 172,281 | 72,207 | 44,543 | 10,988 |
| | Homostylous | *L. bienne* | 73.9 | 462.0x | 385,165 | Circular* | 153,096 | 81,759 | 32,117 | 10,951 |
| | | *L. decumbens* | - | - | - | - | 157,739 | 80,618 | 32,321 | 10,458 |
| | | *L. leonii* | 120.5 | 492x | 282,740 | Circular | 173,198 | 72,468 | 44,895 | 10,940 |
| | | *L. lewisii* | - | - | - | - | 159,778 | 83,267 | 32,810 | 10,891 |
| Clade B2 | Distylous | *L. campanulatum* | 123.9 | 505.8x | 259,104 | Circular | 153,941 | 83,701 | 25,768 | 18,704 |
| | | *L. capitatum* | 122.8 | 501.2x | 255,813 | Circular | 153,941 | 83,701 | 25,768 | 18,704 |
| | | *L. flavum* | 122.4 | 499.4x | 252,293 | Circular | 153,940 | 83,698 | 25,768 | 18,706 |
| | | *L. maritimum* | 115.4 | 471.0x | 241,529 | Circular | 153,917 | 83,543 | 25,642 | 19,090 |
| | | *L. suffricatosum* | 107.5 | 438.8x | 226,675 | Circular | 153,455 | 83,514 | 25,592 | 18,757 |
| | | *L. tauricum* | 121.8 | 496.9x | 254,138 | Circular | 153,828 | 83,586 | 25,768 | 18,706 |
| | | *L. tenue* | 119.3 | 486.7x | 249,133 | Circular | 153,441 | 83,393 | 25,646 | 18,756 |
| | | *L. thracicum* | 113.1 | 461.6x | 236,761 | Circular | 153,941 | 83,701 | 25,768 | 18,704 |
| | Homostylous | *L. teniufolium* | 125.5 | 512.1x | 261,892 | Circular | 153,393 | 83,499 | 25,578 | 18,738 |
| | | *L. trigynum* | 122.4 | 499.4x | 255,131 | Circular | 153,257 | 83,147 | 25,603 | 18,904 |
| Outgroup | Distylous | *T. sinensis* | - | - | - | - | 154,702 | 84,141 | 25,827 | 18,907 |
| Mean | | | 117.51 | 490.45 | 264,160 | - | 157,405 | 81,612 | 29,659 | 14,796 |

**Cov**: coverage; **Asm**: assembly; **Graph**: shape of the graph produced by both GetOrganelle and GeSeq; **LSC**: Large Single Copy region; **IR**: Inverted Repeat; **SSC**: Small Single Copy region; **K115**: the size of the k-mer used to construct the assembly and the graph; **"*":** for this species, the k-mer size was of 85 and not 115; **"-":** data from NCBI so the assembly was already available and we just conducted annotation for these species.

**Suppl. Table S3 –** Hypotheses tested using PAML-codeml, Branch Model

| | |
|---|---|
| **H0** | One dN/dS for the whole tree |
| **H1** | One dN/dS per mating-system |
| **H2** | One dN/dS per mating-system and per clade |

**Suppl. Table S4 –** Log likelihood-ratio test results for D1 nuclear genes

| **D1, nuclear tree** | **LRT** | **DDL** | **PVALUES** |
|---|---|---|---|
| Whole VS mating-system | 30.90 | 2 | <0.001 |
| Mating-system VS Clade + mating-system | 6.31 | 2 | 0.04 |

**Suppl. Table S5** – dN/dS values for nuclear datasets D1 estimated using PAML – codeml, H2 branch-model

| | **CLADE A** | **CLADE B2** |
|---|---|---|
| **Distylous** | 0.10 ± 0.003 | 0.11 ± 0.003 |
| **Homostylous** | 0.14 ± 0.008 | 0.12 ± 0.01 |

**Suppl. Table S6 – Genome assembly statistics**

| **Metric** | **hifiasm+hic primary** |
|---|---:|
| Number of contigs | 850 |
| Largest contig | 54,108,257 |
| Total length* | 888,529,764 |
| GC (%) | 41.12 |
| N50** | 21,077,493 |
| N75 | 9,821,036 |
| L50*** | 12 |
| L75 | 28 |
| # N's per 100 kbp | 0 |
| BUSCO | 93.9% |

*Flow-cytometry based genome size estimates for *L. leonii* is 983 Mb
**N50: 50% of contigs in the assembly are longer than the N50 length; N75: 75%
***L50: Count of the smallest number of contigs which together make up 50% of the genome assembly size; L75: same but up 75%

**Suppl. Table S7 - TE composition of the *L. leonii* genome assembly.**

| Repeats Category | Number of elements | Length (bp) | Percentage of sequence |
|---|---|---|---|
| **Retroelements** | 490,427 | 520,484,188 | 58.6 |
| SINEs | 2,069 | 316,608 | 0.04 |
| LINEs | 31,152 | 28,943,163 | 3.26 |
| RTE/Bov-B | 591 | 58,976 | 0.01 |
| L1/CIN4 | 30,348 | 28,840,452 | 3.25 |
| LTR elements | 457,206 | 491,224,417 | 55.3 |
| Ty1/Copia | 82,672 | 40,299,602 | 4.54 |
| Ty3/DIRS1 | 372,632 | 450,485,038 | 50.72 |
| Retroviral | 1,902 | 439,777 | 0.05 |
| **DNA transposons** | 264,662 | 99,233,497 | 11.17 |
| hobo-Activator | 47,199 | 12,779,374 | 1.44 |
| Tc1-IS630-Pogo | 26,786 | 6,103,395 | 0.69 |
| Tourist/Harbinger | 35,897 | 10,476,496 | 1.18 |
| Other (Mirage/P-element/Transib) | 113 | 22,172 | 0 |
| **Rolling-circles** | 46,008 | 19,847,558 | 2.23 |
| **Unclassified** | 18,877 | 10,154,143 | 1.14 |
| **Total interspersed repeats** | - | 629,871,828 | 70.91 |
| **Small RNA** | 2,069 | 316,608 | 0.04 |
| **Simple repeats** | 73,815 | 3,199,627 | 0.36 |
| **Low complexity** | 11,88 | 599,02 | 0.07 |

Total length = 888,247,830 bp ; GC level = 41.12%

**Suppl. Table S8 – Sample information for *L. leonii* and *L. perenne* population genomic analyses**

| Species | accession/origin | Lat | Lon | Nb of inds analyzed | Nb of downsampled inds |
|---|---|---|---|---|---|
| *Linum perenne* | ger2 near Othfresen, Germany | 52,01 | 10,40 | 63 | 37 |
| | ger3 near Marienhagen, Germany | 52,04 | 9,68 | 54 | 33 |
| | ger5 near Goslar, Germany | 51,90 | 10,52 | 20 | 13 |
| *Linum leonii* | ger4 near Einbeck, Germany | 51,82 | 9,91 | 20 | 10 |
| *Linum leonii* | near Moringen, Germany | 51,73 | 9,80 | 1 (assembly) | |
| *Linum grandiflorum* | *Lgra-62-06-thrum; LIN 10 (IPK)* | NA | NA | 1 | |

**Suppl. Table S9 – Read mapping information**

| Analysis | Species' reads | Species' assembly for read mapping |
|---|---|---|
| **Inbreeding coefficient** | *L. leonii* | *L. leonii* |
| | *L. perenne* | *L. perenne* |
| **Genome-wide $\pi$** | *L. leonii* | *L. leonii* |
| | *L. perenne* | *L. perenne* |
| **Tajima's D** | *L. leonii* | *L. leonii* |
| | *L. perenne* | *L. perenne* |
| **piN/piS** | *L. leonii* | *L. leonii* |
| | *L. perenne* | *L. perenne* |
| **PCA** | *L. leonii + L. perenne* | *L. perenne* |
| **Admixture** | *L. leonii + L. perenne* | *L. perenne* |
| **Dadi** | *L. leonii + L. perenne* | *L. perenne* |
| **DFE** | *L. leonii + L. grandiflorum* | *L. leonii* |
| | *L. perenne + L. grandiflorum* | *L. perenne* |

**Suppl. Table S10 – ADMIXTURE Cross-validation values**

| K | CV value |
|---|---|
| K1 | 0.55028 |
| K2 | 0.43650 |
| K3 | 0.37169 |
| K4 | 0.34807 |
| K5 | 0.33965 |
| K6 | 0.33442 |
| K7 | 0.32908 |
| K8 | 0.32303 |
| K9 | 0.32337 |
| K10 | 0.32648 |

#### Supplementary table S11 – *dadi* 2D models' parameters and descriptions.

| Pop. 1 | Pop. 2 | Best inferred model | Model parameters | | | | | | | | | | | | Year-scaled parameters (generations * 2) | |
|---|---|---|---|---|---|---|---|---|---|---|---|---|---|---|---|---|
| | | | theta | L | Ne_ref | Tsplit | N1 | N2 | T change | N1 change | N2 change | F1 | F2 | AIC[1] | Tsplit scaled | Tchange scaled |
| L.per - ger2 | Lleo | no_mig_size | 2,715 | 66,186 | 1,444,645 | 897,528 | 4,328,838 | 3,036,232 | 62,356 | 33,871 | 11,352 | 0.01 | 0.20 | 4,348 | 1,795,056 | 124,713 |
| Lper - ger3 | Lleo | split_growth | 3,186 | 66,116 | 1,696,929 | 496,799 | 115,634 | 18,129 | | | | 0.00 | 0.55 | 4,57 | 993,598 | |
| Lper - ger5 | Lleo | split_growth | 3,059 | 65,424 | 1,646,582 | 973,122 | 291,993 | 41,574 | | | | 0.00 | 0.21 | 3,325 | 1,946,244 | |

**Model description:**
- *split_growth* (Suppl. Fig. S5-A): Split at Tsplit; two exponential growing populations; inbreeding ($F_1$, $F_2$).
- *no_mig_size* (Suppl. Fig. S5-D): Split at Tsplit; after split, each daughter population has constant size (N1, N2); then at time Tchange, both change to sizes (N1-change, N2-change); inbreeding ($F_1$, $F_2$).

[1]AIC scores are comparable among different tested scenarios for the same population pairs but not between population pairs. Only the score for best-supported demographic history model is presented in this table.

#### Suppl. Table S12 – Hybpiper statistics with number of mapped reads and genes with paralog warning for each sample.

| | Species | Mating system | Number of reads from target capture | | Number of gene with paralog warning | |
|---|---|---|---|---|---|---|
| | | | Total | Mapped | On gene's length | On reads depth |
| **Clade A** | *L. austriacum* | Distylous | 20,228,044 | 6,972,245 | 8 | 9 |
| | *L. grandiflorum* | | 216,368,754 | 178,416 | 11 | 21 |
| | *L. hirsutum* | | 17,055,766 | 2,806,819 | 9 | 15 |
| | *L. narbonense* | | 19,592,244 | 2,154,900 | 16 | 34 |
| | *L. perenne* | | 16,814,040 | 3,028,139 | 15 | 21 |
| | *L. bienne* | Homostylous | 104,067,116 | 118,560 | 9 | 31 |
| | *L. decumbens* | | 83,925,414 | 39,636 | 1 | 2 |
| | *L. leonii* | | 18,208,918 | 5,236,548 | 11 | 18 |
| | *L. lewisii* | | 84,213,206 | 49,799 | 2 | 3 |
| **Clade B2** | *L. campanulatum* | Distylous | 24,272,000 | 4,303,828 | 18 | 34 |
| | *L. capitatum* | | 15,451,632 | 2,649,107 | 14 | 28 |
| | *L. flavum* | | 15,987,986 | 2,723,333 | 17 | 32 |
| | *L. maritimum* | | 17,128,434 | 1,861,584 | 42 | 78 |
| | *L. suffruticosum* | | 16,395,940 | 4,249,703 | 19 | 73 |
| | *L. tauricum* | | 14,253,776 | 2,421,890 | 14 | 29 |
| | *L. tenue* | | 19,575,804 | 4,972,442 | 43 | 90 |
| | *L. thracicum* | | 18,932,174 | 3,282,495 | 16 | 40 |
| | *L. tenuifolium* | Homostylous | 15,125,490 | 3,576,330 | 6 | 14 |
| | *L. trigynum* | | 21,819,288 | 5,604,946 | 42 | 69 |

### DATASET S1

| BUSCO | |
|---|---|
| C:93.9%[S:88.7%,D:5.2%],F:1.3%,M:4.8%,n:2326 | |
| 2,184 | Complete BUSCOs (C) |
| 2,063 | Complete and single-copy BUSCOs (S) |
| 121 | Complete and duplicated BUSCOs (D) |
| 30 | Fragmented BUSCOs (F) |
| 112 | Missing BUSCOs (M) |
| 2,326 | Total BUSCO groups searched |

| REPEAT MASKING | | | | | | |
|---|---|---|---|---|---|---|
| **Species** | **Tool** | **Library** | **Number** | **Total size (kb)** | **Mean size (bp)** | **% Genome** |
| *Linum leonii* | Repeat masker | De-novo | 722,584 | 665,784.20 | 921.39 | 74.95 |
| *Linum leonii* | Repeat Runner | MAKER TE | 30,091 | 26,258.26 | 872.63 | 2.96 |

| INTERPROSCAN | | | | |
|---|---|---|---|---|
| **Databases** | **Nb term linked to mRNA** | **Nb mRNA with term in raw file** | **Nb mRNA updated by term in our annotation file** | **Nb gene updated by term in our annotation file** |
| AntiFam | 25 | 25 | 25 | 22 |
| CDD | 44,825 | 44,825 | 37,978 | 10,586 |
| Coils | 16,886 | 16,886 | 16,886 | 5,802 |
| FunFam | 62,387 | 62,387 | 38,625 | 10,034 |
| GO | 298,695 | 298,695 | 108,379 | 31,984 |
| Gene3D | 118,148 | 118,148 | 78,358 | 20,971 |
| Hamap | 3,098 | 3,098 | 3,032 | 1,184 |
| InterPro | 361,926 | 361,926 | 108,379 | 31,984 |
| MetaCyc | 21,552,435 | 21,552,435 | 79,135 | 21,527 |
| MobiDBLite | 46,057 | 46,057 | 46,057 | 17,801 |
| PANTHER | 106,857 | 106,857 | 106,857 | 31,456 |
| PIRSF | 4,824 | 4,824 | 4,760 | 1,479 |
| PRINTS | 19,772 | 19,772 | 17,049 | 4,181 |
| Pfam | 145,874 | 145,874 | 100,472 | 28,893 |
| ProSitePatterns | 27,351 | 27,351 | 21,710 | 5,275 |
| ProSiteProfiles | 51,123 | 51,123 | 39,918 | 11,783 |
| Reactome | 75,308,129 | 75,308,129 | 84,213 | 23,235 |

| | | | | |
|---|---|---|---|---|
| SFLD | 2,787 | 2,787 | 1,374 | 328 |
| SMART | 40,428 | 40,428 | 32,187 | 8,232 |
| SUPERFAMILY | 97,098 | 97,098 | 76,603 | 20,527 |
| TIGRFAM | 10,595 | 10,595 | 97,75 | 2,852 |

| **CODING GENES ANNOTATION STATISTICS** | |
|---|---|
| Description *Linum leonii* | |
| Number of protein-coding genes | 45,638 |
| Number of mRNA | 13,2390 |
| Average number of exons per mrna | 5.3 |
| Average exon length | 299 |
| Average intron length | 342 |

### SUPPLEMENTARY NOTES

**<u>Supplementary Note S1:</u> Demographic history inference for 2D SFS between *L. leonii* and populations of *L. perenne* ger3 and ger5.**

To estimate the divergence and subsequent demographic history of both species, we inferred split models between *L. leonii* and the three populations of *L. perenne* (ger2, ger3 and ger5) using *dadi* (Gutenkunst et al. 2009; Blischak et al. 2020) (Suppl. Fig. S5, Suppl. Table S10). Results for the split between *L. leonii* and *L. perenne* – ger2 are reported in the main text. Below you can find the results for the two other splits: *L. leonii* vs *L. perenne* - ger3 and *L. leonii* vs *L. perenne* - ger5. The best-fitting split model between *L. perenne* population ger3 and *L. leonii* was *"split_growth"* and indicated a split around 1 Mya. After divergence, both populations experienced exponential size reductions. Population *L. perenne* - ger3 has a final effective population size of ~115,000, while *L. leonii* stabilized at a much smaller effective size of ~18,000. Inbreeding was low in *L. perenne* - ger3 ($F_{ger3} < 0.01$) but higher in *L. leonii* ($F_{leonii\text{-}ger3} = 0.55$) (Suppl. Table S11). These results might be affected by a small amount of gene flow between these two populations which was inferred by ADMIXTURE (Alexander et al. 2009) and was not included in the demographic history models. The same model was preferred for the pair of *L. perenne* - ger5 and *L. leonii*. The split was inferred at ~2Mya and the final effective populations sizes were estimated at ~290,000 and ~40,000 for *L. perenne* - ger5 and *L. leonii* respectively. Inbreeding was similar to the *L. perenne* - ger2 vs *L. leonii* pair ($F_{ger5} < 0.01$, $F_{leonii\text{-}ger5} = 0.21$) (Suppl. Table S10).

Supplementary Note S2: **fastDFE best supported models**

The Distribution of Fitness Effect (DFE) was inferred using fastDFE (Sendrowski and Bataillon 2024) for each population. This software uses and transforms the neutral site frequency spectrum (SFS) to correct for the effect of demography and applies the same transformation to the selected SFS. The selected SFS is then compared to expectations under different distributions of fitness effects to estimate the parameters that best explain the data. The best supported models inferred combined gamma (for deleterious mutations) and exponential (for beneficial mutations) distributions (GammaExpParametrization()) for the three *L. perenne* populations' DFEs, and gamma deleterious-only distribution (using GammaExpParametrization() or GammaDiscreteParametrization()) for *L. leonii*.
